# A hidden T-DNA-linked inversion-duplication causes a pronounced light-dependent phenotype in Arabidopsis

**DOI:** 10.64898/2026.03.19.712841

**Authors:** Maria del Pilar Martinez, Julie Anne V. S. de Oliveira, Ioana Nica, Noah Ditz, Ke Zheng, Vera Wewer, Sabine Metzger, Philipp Westhoff, Holger Eubel, Iris Finkemeier, Markus Schwarzländer, Boas Pucker, Veronica G. Maurino

**Affiliations:** Molecular Plant Physiology, Institute for Cellular and Molecular Botany (IZMB), University of Bonn, Kirschallee 1, 53115 Bonn, Germany; Plant Biotechnology and Bioinformatics, Institute for Cellular and Molecular Botany (IZMB), University of Bonn, Kirschallee 1, 53115 Bonn, Germany; Institute of Plant Genetics, Leibniz University Hannover, Herrenhäuser Str. 2, 30419 Hannover, Germany; Plant Energy Biology, Institute of Plant Biology and Biotechnology (IBBP), University of Münster, Schlossplatz 8, 48143 Münster, Germany; Plant Metabolism and Metabolomics Facility, Institute for Plant Sciences, University of Cologne and Cluster of Excellence on Plant Sciences (CEPLAS), Zülpicher Strasse 47b, 50674 Cologne, Germany; Plant Metabolism and Metabolomic Facility, Heinrich Heine University Düsseldorf and Cluster of Excellence on Plant Sciences (CEPLAS), Universitätsstraße 1, 40225 Düsseldorf, Germany; Plant Physiology, Institute of Plant Biology and Biotechnology (IBBP), University of Münster, Schlossplatz 7, 48149 Münster, Germany

**Keywords:** gene dosage, genotype-phenotype interpretation, large scale chromosomal rearrangement, T-DNA insertion lines

## Abstract

T-DNA insertion mutants are widely used to disrupt genes and infer their functions, yet the insertions can also trigger unintended genomic changes that confound phenotypic interpretation. Here, we used T-DNA insertion mutants affecting the major mitochondrial malate dehydrogenase (MDH1) and the heterodimeric NAD-dependent malic enzymes (ME1 and ME2) to examine their functional coordination across photoperiods and irradiance regimes. Under short days, especially at low light intensity, *mdh1xme2* mutants were markedly smaller than wild type and, unexpectedly, than the *mdh1xme1xme2* triple mutant, and they showed a more pronounced reduction in photosynthetic capacity. ME1 was undetectable in *mdh1xme2*, implying that the double and triple mutants effectively lack heterodimeric ME and should therefore behave similarly, contrary to what we observed. Whole-genome analysis resolved this discrepancy by revealing that the *MDH1* T-DNA insertion in *mdh1xme2* is accompanied by a major rearrangement, a 137-kbp duplication downstream of the insertion site, which was absent in the *mdh1xme1xme2* triple mutant. This duplication increased gene dosage and elevated transcript abundance across the duplicated interval, while proteomics detected 5 of the 38 encoded proteins, including PEPC1. *mdh1xme2* accumulated oxaloacetate-derived amino acids and displayed an altered carbon/nitrogen balance, making PEPC1 a plausible contributor to the exacerbated *mdh1xme2* phenotype. Together, our data indicate that a T-DNA-linked structural variant can amplify expression of dozens of genes and intensify phenotypes at specific conditions, thereby affecting the interpretation of genotype-phenotype relationships. Because *Agrobacterium*-mediated DNA transfer also underpins many genome-editing workflows, our findings argue that structural validation around insertion/editing loci should be considered essential when interpreting T-DNA-derived plant lines.

## Introduction

*Agrobacterium tumefaciens* has been a cornerstone of plant genetic transformation for decades, enabling efficient delivery of DNA into the nuclear genome and thereby accelerating functional genomics in many species. In its natural infection process, *A. tumefaciens* transfers a defined segment of its tumor-inducing (Ti) plasmid, termed transferred DNA (T-DNA), into plant cells, where it integrates into the host genome and drives crown gall formation (Van Larebeke et al., 1975; Chilton et al., 1977). T-DNA transfer relies on virulence (*vir*) genes located on the Ti plasmid but outside the T-DNA region (Akiyoshi et al., 1984; Schröder et al., 1984; Zambryski et al., 1989). The T-DNA itself is delimited by imperfect ∼25-bp border repeats (left and right borders; LB and RB) that are required for processing and integration. The internal native T-DNA sequence can be replaced by engineered cargo while retaining the borders, which forms the very basis of routine plant transformation (Wang et al., 1984). The development of binary vector systems, in which the T-DNA and *vir* functions are provided on separate replicons, further simplified transformation workflows and enabled large-scale mutant generation (de Framond et al., 1983; Hoekema et al., 1983; Alonso et al., 2003; Lee and Gelvin, 2008; Szarzanowicz et al., 2025).

Beyond transgenesis, T-DNA insertion mutagenesis has become a standard strategy to infer gene function by disrupting coding sequences or regulatory regions (Marks and Feldmann, 1989; Koncz et al., 1990; Feldmann, 1991). Following publication of the first *Arabidopsis thaliana* genome sequence (Arabidopsis Genome Initiative, 2000), community resources expanded rapidly. Indexed insertion collections, together with searchable databases, enabled routine identification of presumed loss-of-function alleles (Meinke et al., 2003; Alonso and Ecker, 2006). Today, hundreds of thousands of *A. thaliana* insertion lines are available, including major collections such as SALK, GABI-Kat, SAIL, and WISC (Sussman et al., 2000; Sessions et al., 2002; Alonso et al., 2003; Rosso et al., 2003; Kleinboelting et al., 2012; O’Malley et al., 2015). This scale has made T-DNA lines indispensable for dissecting pathways ranging from metabolism to development and stress signaling.

Yet, the apparent genetic simplicity insertion of T-DNAs into plant genomes has limits that may be overlooked. While T-DNA integration is frequently described as largely random and stable (Thomas et al., 1994; Azpiroz-Leehan and Feldmann, 1997; Gelvin, 2017), its molecular route is closely tied to host DNA double-strand break repair. In practice, many insertion events are complex: truncated borders, tandem repeats, multiple T-DNA copies per genome (with an average of 1.5 T-DNA copy per T-DNA line) and the inclusion of unintended plasmid backbone sequence are common (Castle et al., 1993; Gelvin, 2003; O’Malley et al., 2015; Gelvin, 2017; Nicolia et al., 2017; Pucker et al., 2021). Moreover, DSB repair itself can generate structural variants, via processes such as single-strand annealing or non-allelic homologous recombination. Also, T-DNA lines have repeatedly been found to carry large deletions, inversions, duplications, insertions, and translocations linked to the integration event (Nacry et al., 1998; Clark and Krysan, 2010; Krispil et al., 2020). Such collateral changes can alter gene dosage or regulatory landscapes and may confound genotype-phenotype interpretation, especially when phenotypes are subtle, conditional, or interpreted through genetic comparisons (Tax and Vernon, 2001; Valentine et al., 2012; Tamura et al., 2016). To characterize T-DNA insertion lines, several techniques can be used, ranging from PCR-based approaches to long-read genotyping and digital droplet PCR (Liu et al., 1995; O’Malley et al., 2007; Głowacka et al., 2016; Pucker et al., 2021).

In attempt to elucidate how mitochondrial malate metabolism affects plant development and physiological performance across different light environments, we examined *A. thaliana* T-DNA insertion lines impaired in the major mitochondrial malate dehydrogenase (MDH1) and the heterodimeric NAD-dependent malic enzyme (ME1 and ME2). Interestingly, the *mdh1.1xme2* double mutant exhibited a much stronger light-dependent phenotype than the *mdh1.1xme1xme2* triple mutant, suggesting an additional layer of genetic complexity beyond the intended mutations. Here, we shed light on the genetic basis of this observation by combining physiological phenotyping with whole-genome analysis. Our data illustrates how insertion-associated genome restructuring can influence mutant phenotypes. By linking a T-DNA-associated large-scale duplication to altered gene dosage and downstream molecular and physiological readouts, our study provides a compelling case for the need of caution when inferring functional effects based on T-DNA-insertions, validating genome structure around insertion alleles emerges as an important prerequisite for attribution of causality.

## Results

### Unexpectedly exacerbated phenotype of *mdh1.1xme2* relative to *mdh1.1xme1xme2* under short days

Recently, using T-DNA insertion mutants lacking the major mitochondrial malate dehydrogenase (MDH1) and the two subunits of NAD-malic enzyme (ME1 and ME2), we identified mitochondrial malate decarboxylation capacity as a key control point linking respiratory energy supply to plastid function under low-energy conditions (Martinez et al., 2026). Because NAD-ME acts predominantly as a heterodimer *in vivo* (Tronconi et al., 2008), we hypothesized that if leaf NAD-ME function depends on heterodimer assembly, then double mutants combining *mdh1.1* with loss of the single ME2 subunit (*mdh1.1xme2.1* and *mdh1.1xme2.2;* Suppl. Fig. 1; Suppl. Table 1) should phenocopy the *mdh1.1xme1xme2* triple mutant.

To test this hypothesis, we analyzed growth of all genotypes across photoperiods and light intensities. Under long-day conditions (LD) at 120 µmol photons m^-2^ s^-1^ (normal light, NL) we observed no visible differences among genotypes (Fig. 1). In striking contrast, under short-day (SD) conditions the double mutants exhibited a pronounced phenotype: they were pale green and severely growth impaired relative to wild type (Fig. 1; Suppl. Fig. 1). Unexpectedly, this defect was stronger than in the triple mutant (Fig. 1; Suppl. Fig. 1). In SD at 60 µmol photons m^-2^ s^-1^ (low light, LL), the triple mutant remained smaller than wild type, but it was consistently larger than both double-mutants. In SD and NL, the triple mutant was largely indistinguishable from the wild type, whereas the double mutants remained markedly stunted. Moreover, the double mutants did not flower in SD. Together, these observations suggest that the *mdh1.1xme2* double mutants carry an additional factor that exacerbates growth defects, particularly under low-energy conditions.

**Figure 1.**
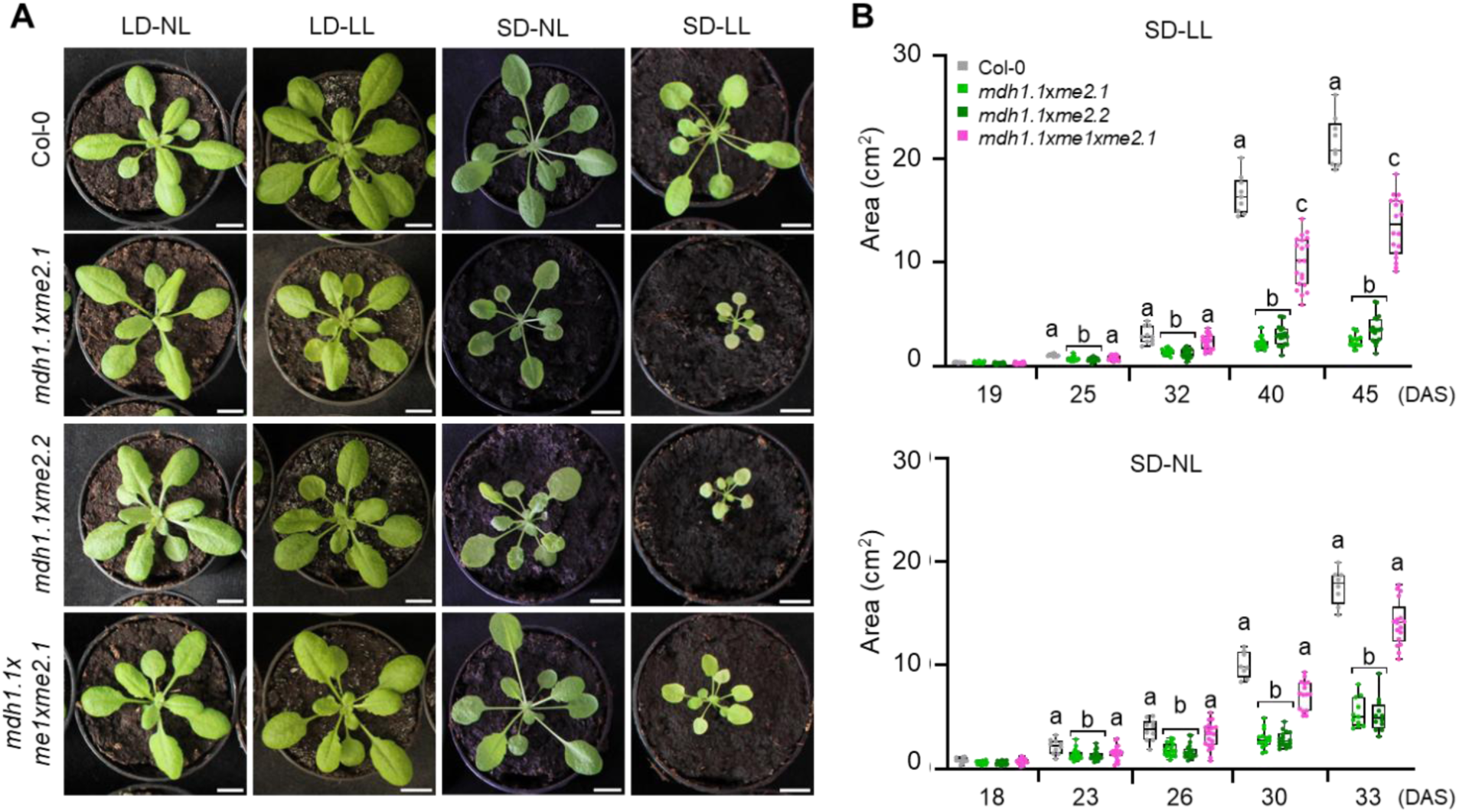
Light-intensity- and photoperiod-dependent phenotypes of plants lacking MDH1 and ME. **A)** Representative plants of each genotype grown under long-day (LD) conditions at 120 µmol photons m^-2^ s^-1^ (NL) for 23 days, short-day (SD) at NL for 30 days, and SD at 60 µmol photons m^-2^ s^-1^ (LL) for 40 days. Scale bar, 1 cm. **B)** Rosette area at different days after sowing (DAS) under SD at NL and LL (n = 10). Statistics were performed separately for each day using one-way ANOVA followed by Tukey’s post-hoc test. Different letters indicate statistically significant differences (*p* < 0.05).

### Low photosynthetic performance and thylakoid remodeling in *mdh1.1xme2*

We assessed the photosynthetic performance of plants grown under SD and exposed to either LL or NL conditions. Under both irradiance regimes, *mdh1.1xme2* exhibited a markedly lower photosynthetic capacity than Col-0 and the triple mutant (Fig. 2A and B). In LL, the maximum quantum efficiency of PSII (Fv/Fm) in *mdh1.1xme2* was reduced to ∼50 % of the value observed under NL. Consistent with strong impairment of PSII function, the effective quantum yield of PSII (ΦPSII) and the electron transport rate (ETR) were close to zero. This disruption also affected energy dissipation, as non-photochemical quenching (NPQ) increased and, in LL reached values ∼2-times higher than in NL. In comparison, the triple mutant also showed reduced photosynthetic parameter values in LL than in NL, but these remained intermediate between those of wild type and double mutants. Under NL, the triple mutant showed wild-type-like values for all measured parameters.

**Figure 2.**
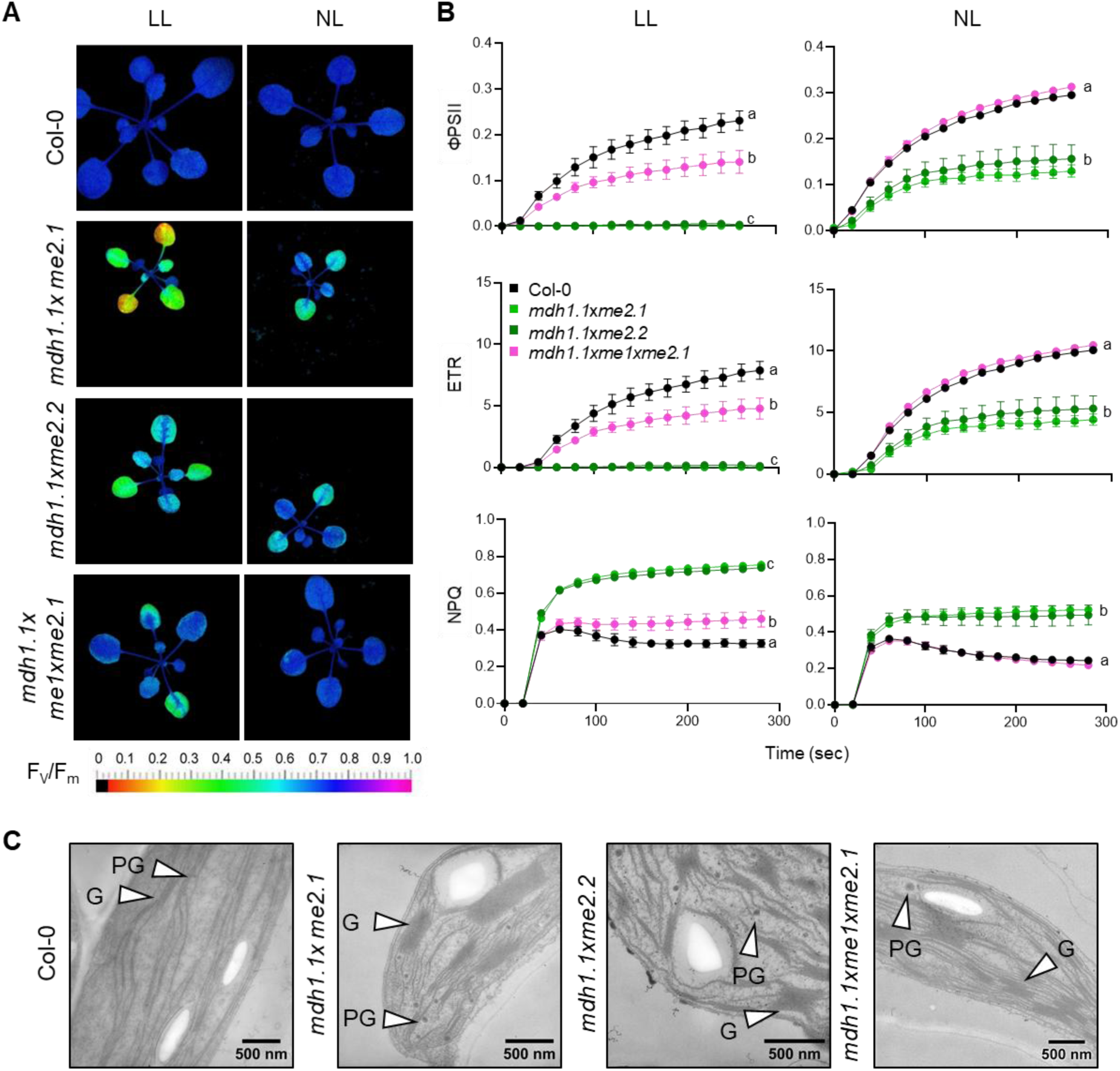
Photosynthetic performance and chloroplast ultrastructure under SD. **A)** Representative false-color images of the maximum quantum yield of PSII (Fv/Fm). **B)** Chlorophyll fluorescence parameters derived from imaging in (A): ΦPSII (effective quantum yield of PSII), ETR (electron transport rate), and NPQ (non-photochemical quenching). Values are means ± SE (n = 3). **C)** Transmission electron micrographs of leaf sections from 40-day-old plants grown under SD at 60 µE and harvested at the end of the night. G, grana; PG, plastoglobules. Data from *mdh1.1xme1xme2.1* were taken from Martinez et al. (2026).

Given the pale green phenotype and reduced photosynthetic capacity of *mdh1.1xme2* mutants, we examined the chloroplast ultrastructure. Under SD-LL conditions, chloroplasts from the double mutant showed marked grana hyperstacking, with thicker and more extensively stacked grana thylakoids, and exhibited an increased abundance of plastoglobules compared with the wild type (Fig. 2C). Enhanced grana stacking is commonly associated with a shift toward higher PSII and LHCII content relative to PSI, and may therefore reflect changes in the composition of photosynthetic complexes and/or pigment complement (Olive and Vallon, 1991; Häusler et al., 2009; van Wijk and Kessler, 2017). Increased plastoglobuli abundance is linked to leaf senescence and stress, consistent with a role for plastoglobules in sequestering lipids and chlorophyll released during thylakoid remodeling (van Wijk and Kessler, 2017; Zechmann, 2019). A less pronounced increase in grana stacking was reported for *mdh1.1xme1xme2*, although without a detectable increase in plastoglobules (Fig. 2C).

### Phytohormone homeostasis is perturbed in *mdh1.1xme2* under SD

Because the double mutant exhibited a more severe phenotype than the triple mutant, we quantified phytohormone profiles in plants grown under SD in LL and NL (Fig. 3). Under both irradiance regimes, *mdh1.1xme2* showed a marked reduction in the growth-promoting auxin indole-3-acetic acid (IAA), which reached only about 50 % of the wild-type level. Salicylic acid (SA) was likewise reduced under both LL and NL to approximately 40% of wild-type levels, arguing against a classical SA-mediated growth-defense trade-off as the primary cause of the growth defect. Instead, reduced SA may reflect limited precursor availability or altered chloroplast metabolism, given the close connection between SA biosynthesis and plastidial function (Roussin-Léveillée et al., 2025). By contrast, jasmonic acid (JA), a stress-associated hormone, accumulated specifically under LL in *mdh1.1xme2*, likely reflecting stronger metabolic imbalance and/or activation of stress-acclimation pathways under these conditions. *cis,trans-*Absicic acid (c,t-ABA) levels in the mutant do not differ from wild type.

**Figure 3.**
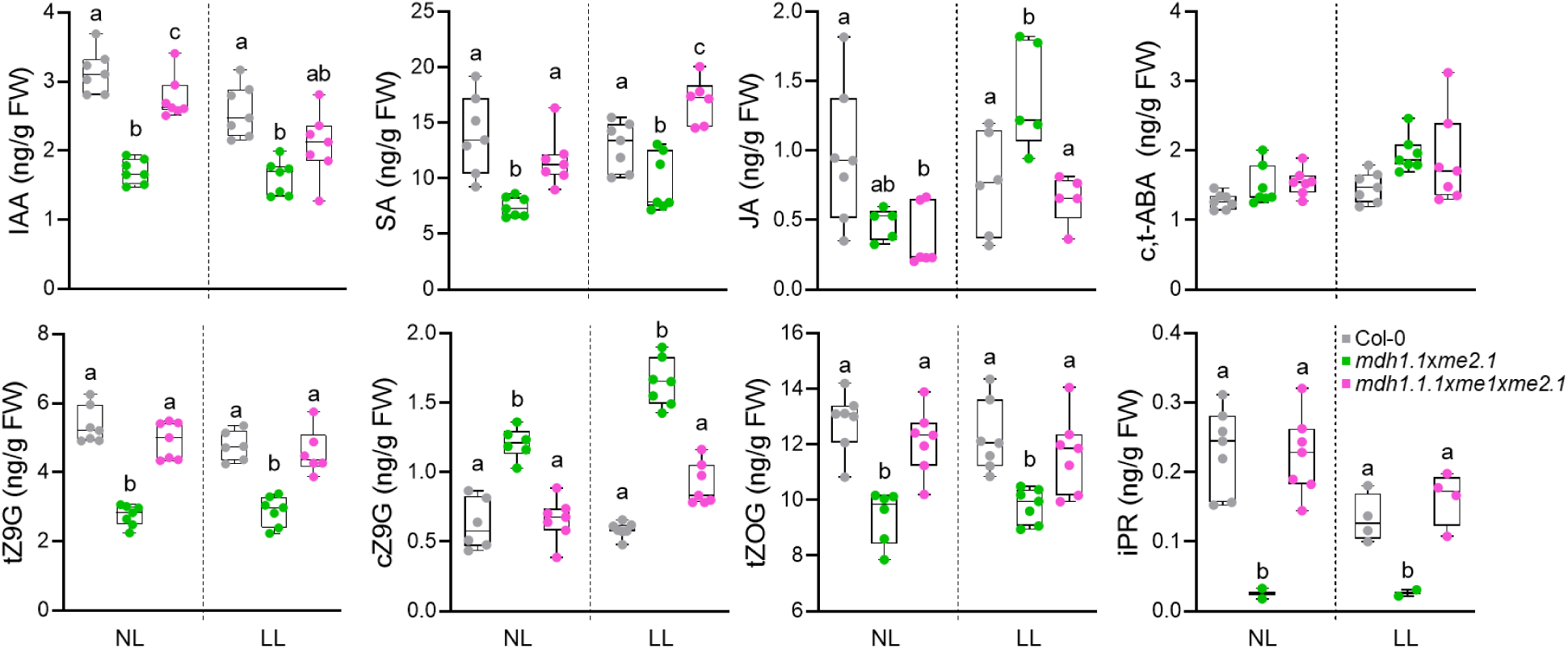
Phytohormone profiling. Whole rosettes were harvested 1 h before lights-on (-1 h) from plants grown under short-day (SD) conditions at normal light (NL) or low light (LL) to quantify phytohormone levels in *mdh1.1xme2.1*, *mdh1.1xme1xme2.1* and Col-0. For each condition, 6-7 independent biological replicates were analyzed, and hormone abundance was normalized to fresh weight. Statistical analyses were performed separately for each light intensity using one-way ANOVA followed by Tukey’s post-hoc test; different letters indicate significant differences (*p* < 0.05).

Cytokinin homeostasis was also substantially disturbed. The trans-zeatin conjugates *trans*-zeatin-9-glucoside (tZ9G) and *trans*-zeatin-O-glucoside (tZOG) were reduced under both irradiance regimes to approximately 50% and 30% of wild-type levels, respectively. In contrast, *cis*-zeatin-9-glucoside (cZ9G) accumulated strongly, reaching about 2.5-times higher levels than in the wild type under NL and roughly 3-fold higher levels under LL. Because *trans*-zeatin-type cytokinins are linked to cell division, meristem function, and leaf and chloroplast development (Dello Ioio et al., 2007; Cortleven and Schmülling, 2015), depletion of the tZ branch is consistent with the reduced growth and impaired photosynthetic performance of *mdh1.1xme2.1*. Conversely, accumulation of cis-zeatin conjugates is linked to growth arrest and the onset of senescence (Cheng et al., 2020), in line with the elevated plastoglobuli abundance observed in the double mutant. Levels of isopentenyladenosine (iPR) were also altered in *mdh1.1xme2.1*. Because iPR-related tRNA modifications contribute to efficient codon-anticodon pairing, disruption of this pathway has been linked to translational defects and mitochondrial metabolic dysfunction (Cheng et al., 2020). Overall, the decline in tZ conjugates together with the rise in cZ9G points to a shift from trans-zeatin-associated growth promotion toward a *cis*-zeatin-associated growth-arrest and senescence-like state, potentially reinforced by organellar metabolic stress. This altered hormonal landscape may contribute directly to dwarfism and chloroplast remodeling in the double mutant, although it may also partly represent a secondary consequence of the underlying mutation-dependent defects.

### ME1 activity is absent in *mdh1.1xme2*

The principal genetic difference between *mdh1.1xme2* and *mdh1.1xme1xme2* is the additional loss of ME1 in the triple mutant. To test whether differences in ME1 abundance or mitochondrial NAD-ME capacity are responsible for the stronger phenotype, we measured NAD-ME activity in mitochondrial extracts, assessed ME1 protein levels by immunoblotting, and quantified protein abundance by LC-MS/MS-based proteomics.

In wild-type mitochondrial extracts, NAD-ME activity was readily detectable, whereas activity in both double and triple mutants was below the detection limit (Fig. 4A). Consistent with this, immunoblotting detected a clear ME1 band in wild type but no immunoreactive signal in either mutant line (Fig. 4B). LC-MS/MS proteomics identified multiple ME1-, ME2-, and MDH1-derived peptides in wild-type samples, whereas none of these peptides were detected in extracts from either the double or triple mutant (Fig. 5C). In contrast, low levels of MDH2-derived peptides were detected at similar abundance in all genotypes. Together, the concordant absence of NAD-ME activity, ME1 immunoreactivity, and a lack of ME1 peptide detection indicates that ME1 does not accumulate in both mutant backgrounds. Thus, *mdh1.1xme2.1* already lacks detectable ME1, suggesting that mitochondrial malate decarboxylation capacity is compromised to the same extent in *mdh1.1xme2.1* and *mdh1.1xme1xme2.1*. Accordingly, the enhanced phenotype of the double mutants is not the result of residual ME1 activity.

**Figure 4.**
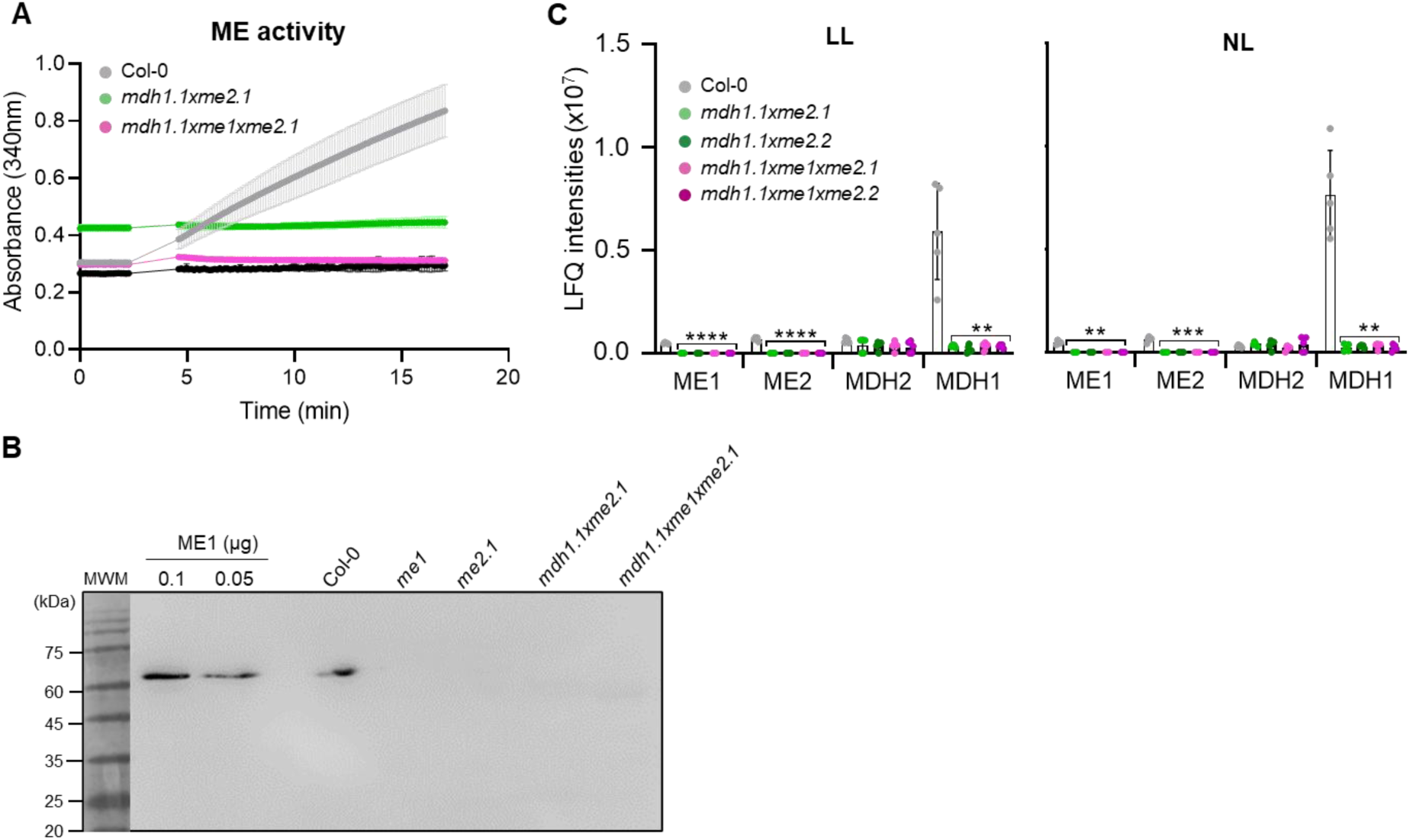
Abundance and activity of mitochondrial ME and MDH in plants grown in SD. **A)** NAD-ME activity in mitochondrial extracts, quantified by monitoring NADH production at 340 nm. Data are means ± SD (n = 3). **B)** Immunoblot of mitochondrial extracts from plants grown under SD at NL and harvested 1 h after lights-on (+1 h), probed with ME1-specific antibodies. 30 µg of mitochondrial proteins were loaded in each well and 7 µl of the MWM. MWM: molecular-weight marker (ROTI®Mark TRICOLOR (Carl Roth)). kDa: kilodaltons. **C)** Label-free quantification (LFQ) of ME1, ME2, MDH1, and MDH2 proteins in plants grown under SD at low light (LL) or normal light (NL). Bars represent means ± SD (n = 5). Significance relative to Col-0 was assessed using Welch’s t-test (p < 0.05, *; p < 0.01, **; p < 0.001, ***; p < 0.0001, ****). Proteomic data from *mdh1.1xme1xme2* was taken from Martinez et al. (2026).

**Figure 5.**
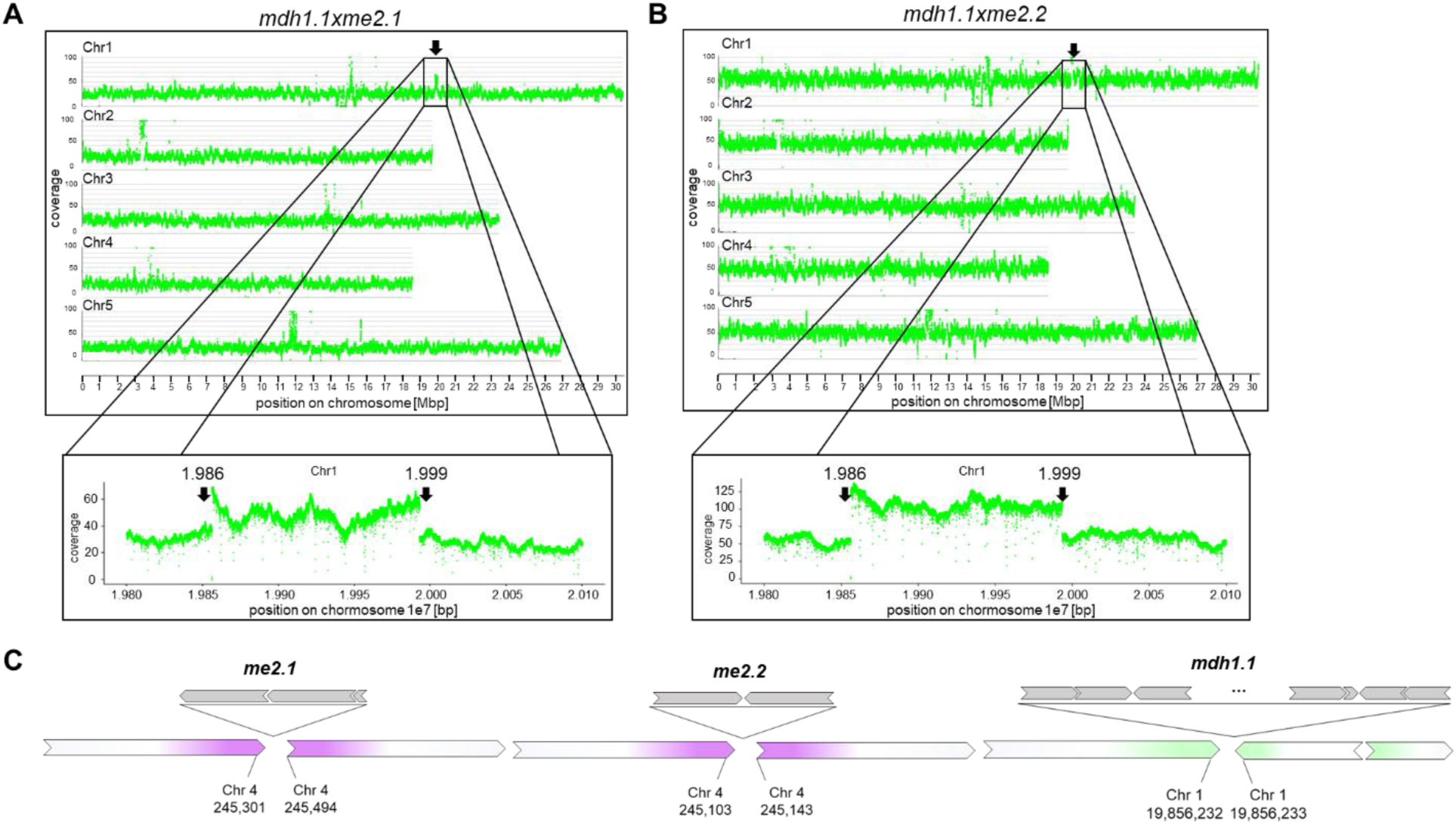
Chromosomal re-arrangements in *mdh1.1xme2* mutant lines. **A-B)** Read-depth coverage plots for (**A**) *mdh1.1xme2.1* and (**B**) *mdh1.1xme2.2*, generated with a custom Python script based on long read mappings (available at GitHub: https://github.com/bpucker/GKseq2). In both double mutants, a duplication was detected downstream of the T-DNA insertion site in *MDH1*. **C)** Schematic overview of the chromosomes carrying the T-DNA insertions. T-DNA insertions are shown in gray, with arrowheads indicating insertion orientation. Segment lengths are not drawn to scale.

### Genome sequencing uncovers a major genomic rearrangement in *mdh1.1xme2*

Given the unexpectedly strong phenotype of *mdh1.1xme2* despite the absence of ME1 protein in this background, we investigated whether additional genomic lesions might contribute to the phenotype. We therefore performed whole-genome sequencing of *mdh1.1xme2.1* and *mdh1.1xme2.2* using Oxford Nanopore Technologies (ONT) long-read sequencing. Local assemblies confirmed the presence of T-DNA arrays at the expected genomic loci, At1g53240 (MDH1) and At4g00570 (ME2) (Fig. 5C).

Because large-scale rearrangements can arise in T-DNA insertion lines (Pucker et al., 2021), we next assessed structural variation by examining read-depth profiles from the ONT datasets. Both double-mutant lines carried a pronounced genomic rearrangement consisting of a ∼137-kb duplication immediately downstream of the *MDH1* T-DNA insertion site. The duplicated segment spans approximately from 1.986 x 10^7^ to 1.999 x 10^7^ bp on chromosome 1 (TAIR10) and is inserted in inverted orientation relative to the reference genome (Fig. 5A, B and C). This duplicated interval encompasses 38 annotated genes (Suppl. Table 1). Interestingly, the duplication was not found in *mdh1.1xme1xme2.1* (Martinez et al., 2026).

### Transcripts encoded in the duplicated region show increased abundance

Following the identification of the genomic rearrangement in both *mdh1.1xme2* mutant lines, we assessed how the duplicated interval affects transcript abundance using RNA-seq. Samples were collected from plants grown under SD conditions in LL and NL, and under LD conditions in NL, at one hour before dawn (-1 h) and one hour after illumination (+1 h). Differential expression was summarized as log_2_ fold-change (log_2_FC) relative to Col-0 and visualized as a heatmap (Fig. 6).

**Figure 6.**
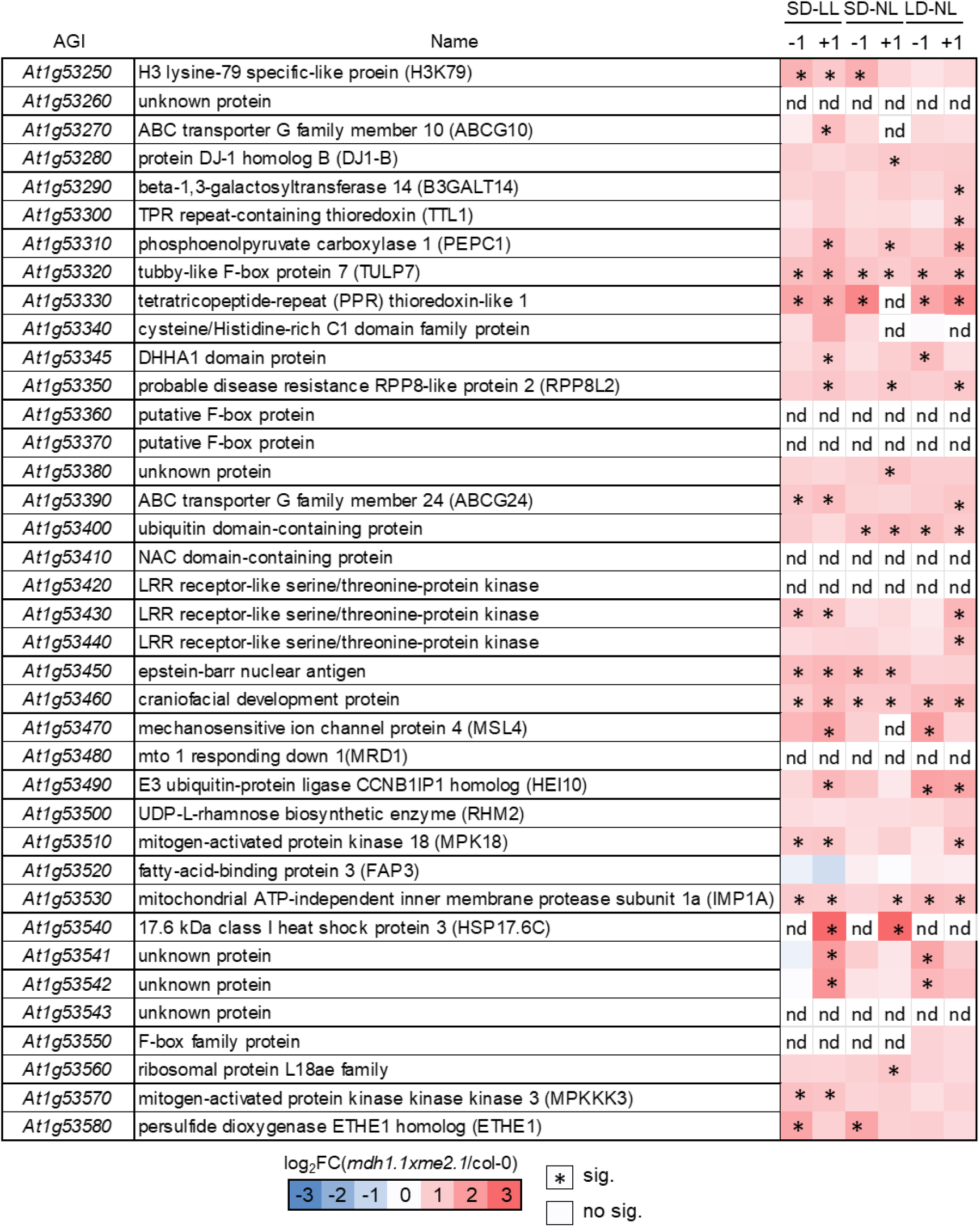
Differential transcript abundance within the duplicated region in *mdh1.1xme2.1*. Gene expression was quantified by RNA-seq in plants grown under short days (SD) at low light (LL) or normal light (NL), under long days (LD) under NL. Samples were harvested 1 h before (-1 h) and 1 h after (+1 h) dawn. Differential expression was summarized as log_2_ fold-change (log_2_FC) relative to Col-0. Transcripts were classified as significantly changed when |log_2_FC| > 1 and padj < 0.05. nd, not detected; sig, significant.

Within the duplicated segment (spanning *At1g53250*-*At1g53580*), transcripts for 32 genes were detected under all analyzed conditions. The largest transcriptional differences between genotypes and irradiance treatments occurred at +1 h. Under SD-LL, 19 of the 32 genes showed significantly higher transcript abundance in the double mutants relative to Col-0 (log_2_FC > 1; padj < 0.05). Under SD-NL, the response was weaker, with 11 genes showing increased transcript levels. At -1 h, 11 genes were already elevated under LL, whereas only 7 transcripts were significantly increased under NL (Fig. 6; Suppl. Data 1).

These results indicate that the duplicated segment drives a light-transition-associated increase in transcript abundance, particularly under LL, suggesting that gene dosage-dependent misexpression contributes to the enhanced, light-dependent phenotype of the duplication-carrying double mutant. To identify genes uniquely or commonly differentially expressed before (-1 h) and after (+1 h) light onset under SD in LL and NL, we generated a Venn diagram (Suppl. Fig. 2; Suppl. Data 1). Below, we focus on differentially expressed genes with functional annotation.

#### Genes commonly accumulated under LL and NL

Three transcripts accumulated across all four conditions (SD-LL -1/+1 and SD-NL -1/+1): *At1g53320*, *At1g53450*, and *At1g53460*. *At1g53320* encodes the F-box protein TULP7, a component of SCF-type E3 ubiquitin ligase complexes involved in proteasome-mediated protein turnover (Gagne et al., 2002; Bashore et al., 2023). Their consistent accumulation suggests a baseline regulatory response around the dark-light transition independent of light intensity. At -1 h, *ETHE1* (*At1g53580*), encoding a mitochondrial sulfur dioxygenase involved in sulfur metabolism (Holdorf et al., 2012), accumulated under both LL and NL. After illumination (+1 h), three transcripts were commonly more abundant under both light intensities: *PEPC1* (*At1g53310*), *RPP8L2* (*At1g53350*), and *HSP17.6C* (*At1g53540*). Upon illumination, *HSP17.6C*, which encodes a small heat-shock protein, was strongly induced (∼32-times under LL and ∼16-times under NL), while *PEPC1*, encoding the anaplerotic enzyme phosphoenolpyruvate carboxylase 1, showed a more moderate increase of approximately threefold under LL and twofold under NL. These results indicate that the night-to-light transition activates metabolic and protein protection mechanisms largely independent of irradiance.

#### Genes specifically accumulated under SD-LL

Several transcripts accumulated preferentially under SD-LL. Four genes showed increased abundance at both time points: *ABCG24* (*At1g53390*), encoding a ABC transporter which are integral membrane protein complexes that use the energy of ATP hydrolysis to transport substrates across membranes; an LRR receptor-like kinase gene (*At1g53430*), predicted to encode a receptor involved in extracellular signal perception; *MAPK18* (*At1g53510*); and *MAPKKK3* (*At1g53570*), both encoding components of MAP kinase signaling pathways. Additional transcript abundances increased after illumination (+1 h), including *ABCG10* (*At1g53270*), predicted to encode an ABC transporter; *MSL4* (*At1g53470*), encoding a mechanosensitive ion channel; and *HEI10* (*At1g53490*), encoding an E3 ubiquitin ligase required for meiosis and overlapping with *MDR1*, a salicylic-acid-responsive gene. Together, these results suggest that higher irradiance preferentially enhances signaling-related responses, including membrane transport and MAP kinase pathways.

#### Genes specifically accumulated under SD-NL

Under SD-NL, *At1g53400*, encoding a ubiquitin-domain protein and putative regulator of ubiquitin-dependent protein degradation, accumulated at both -1 and +1 h. After illumination (+1 h), *At1g53280*, encoding DJ1 homolog B (DJ1B) predicted to function in oxidative stress protection, and *At1g53560*, encoding a structural ribosomal protein of the L18ae family, accumulated specifically under NL conditions. These responses suggest that lower irradiance favors cellular maintenance processes, including redox protection, proteostasis, and translational capacity.

Together, these findings indicate that the duplicated genomic region integrates light-dependent transcriptional responses with pathways involved in metabolism, protein turnover, and stress signaling. While some genes respond similarly to the dark-light transition regardless of irradiance, lower irradiance (LL) primarily activates signaling pathways, whereas normal irradiance (NL) favors maintenance functions related to redox balance and protein homeostasis.

### Duplication-associated accumulation of PEPC1 correlates with the enhanced *mdh1.1xme2* phenotype

To assess duplication-dependent changes in protein abundance, we performed proteome analysis on plants grown under SD in LL and NL and harvested 1.5 h after lights-on. Of the 38 genes encompassed by the duplicated interval, only five corresponding proteins were detected: PEPC1, ETHE1, DJ1B, RHM2 (UDP-L-rhamnose biosynthetic enzyme), and FAP3 (fatty-acid-binding protein 3). RHM2 and FAP3 did not show increased accumulation in any of the conditions analyzed (Fig. 7A; Suppl. Data 2). Under LL, PEPC1, ETHE1, and DJ1B accumulated to higher levels in *mdh1.1xme2* than in Col-0, whereas under NL only PEPC1 and ETHE1 remained significantly increased (Fig. 7A; Suppl. Data 2). By contrast, in *mdh1.1xme1xme2*, only ETHE1 was more abundant under LL, while none of the remaining proteins showed significant increases under either light regime (Fig. 7B; Suppl. Data 2). Accordingly, the increased protein abundance observed in *mdh1.1xme2* is not detected in the triple mutant background lacking the duplicated genomic segment.

**Figure 7.**
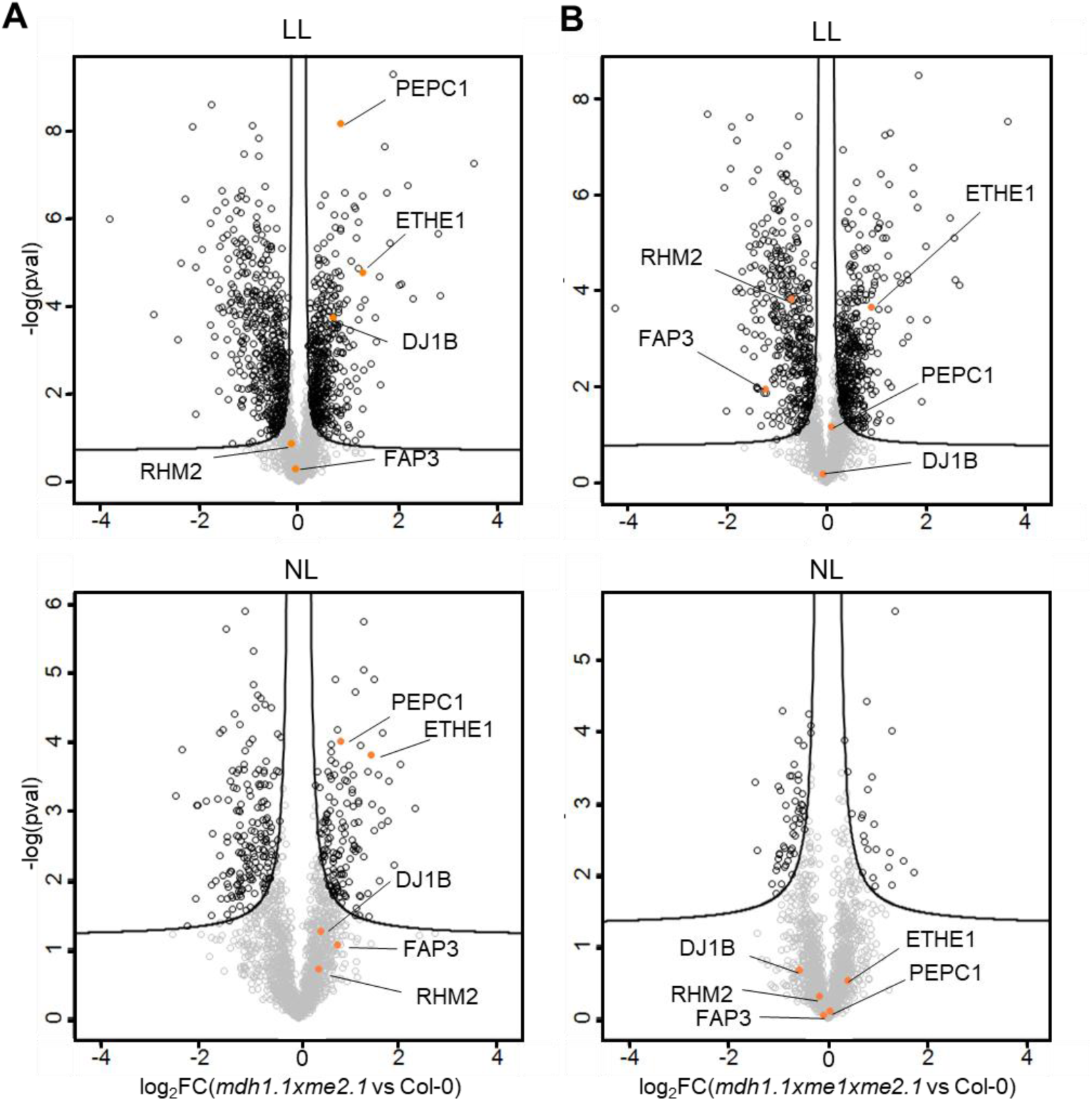
Light-dependent abundance changes in protein groups between *mdh1.1xme2.1, mdh1.1xme1xme2.1*and Col-0 grown under LL and NL in SD. Comparison of log_2_-transformed LFQ-values of (A) *mdh1.1xme2.1* and Col-0, and (B) *mdh1.1xme1xme2.1* and Col-0 protein groups grown in SD at LL or NL employing five independent biological replicates (Suppl. Data 2). Protein groups were graphed by fold change (log_2_FC; x-axis) and the confidence statistic (−log_10_(pvalue); y-axis). Orange points represent the proteins codified by the duplicated region. Protein groups located in the areas above the black lines fulfil the significance threshold limits (FDR ≤ 0.05; S_0_ = 0.1). Data from *mdh1.1xme1xme2.1* was taken from Martinez et al. (2026). ETHE1: ethylmalonic encephalopathy protein1; DJ1B: DJ-1 homolog B; FAP3: fatty-acid-binding protein 3; PEPC1: phosphoenolpyruvate carboxylase 1; RHM2: UDP-L-rhamnose biosynthetic enzyme; LL: low light; NL: normal light; SD: short day.

Consistent with these results, PEPC1 abundance in *mdh1.1xme2* was clearly elevated, while *mdh1.1xme1xme2* displayed PEPC1 levels close to those of the wild type (Fig. 8A). To independently validate the proteomics results, we measured PEPC activity in total protein extracts from plants grown under SD at LL or NL and harvested 1.5 h after lights-on. PEPC activity was significantly higher in *mdh1.1xme2* than in Col-0 under LL (Fig. 8B). We further assessed PEPC1 abundance by immunoblotting using an anti-PEPC1 antibody. A band at ∼110 kDa, consistent with the expected size of PEPC1, was detected with higher intensity in *mdh1.1xme2* than in Col-0 and *mdh1.1xme1xme2*, while both triple mutants displayed band intensities similar to wild type (Fig. 8C).

**Figure 8.**
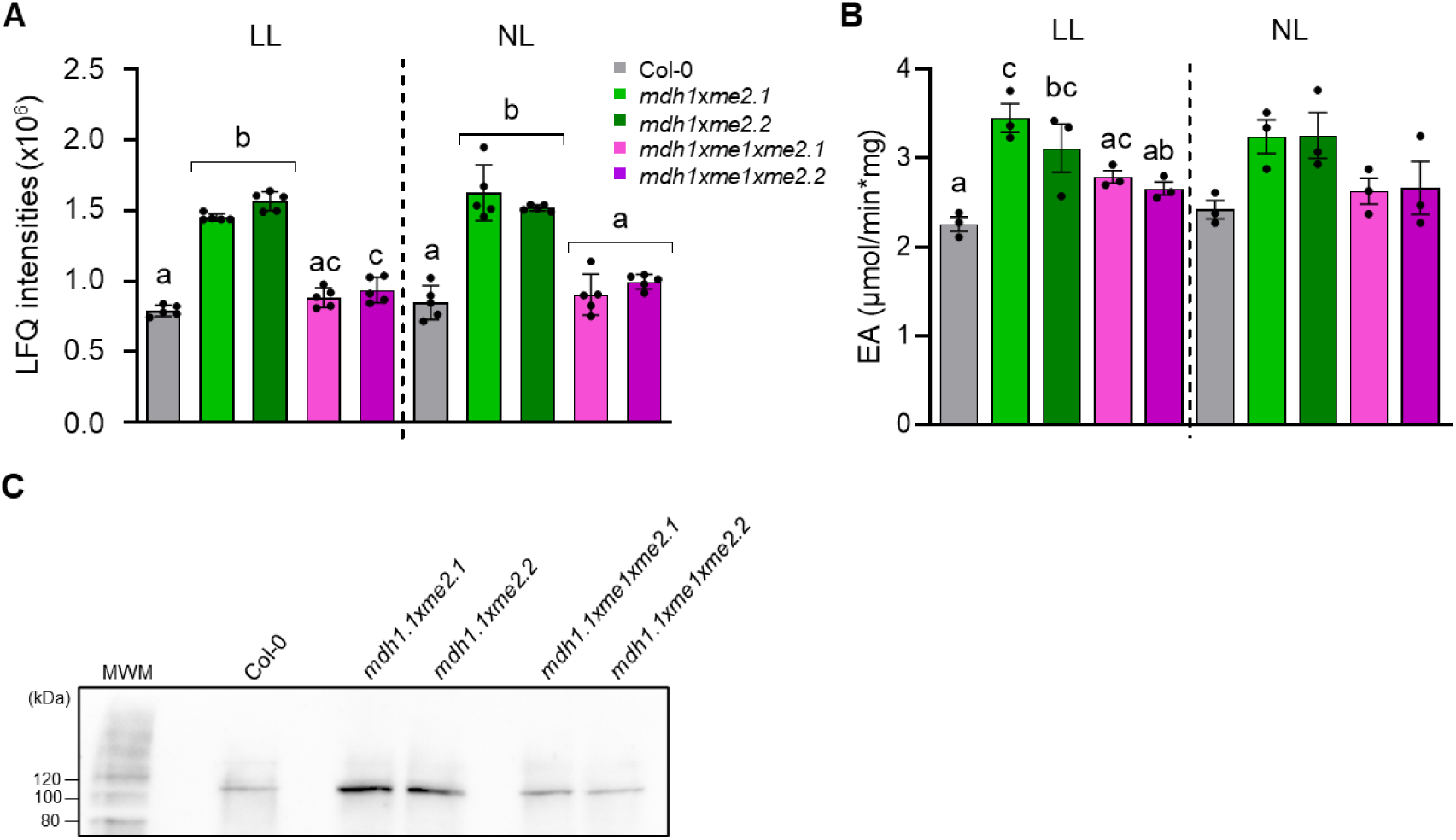
Increased PEPC1 abundance and activity in *mdh1.1xme2*. **A)** Label-free quantification (LFQ) of PEPC1 protein abundance in plants grown under SD at LL or NL. Bars show means ± SD (n = 5). Proteomic data from *mdh1.1xme1xme2.1* was re-analyzed from Martinez et al (2026). **B)** PEPC enzymatic activity in total protein extracts from plants grown under SD at LL or NL, harvested 1 h after lights-on (+1 h). Panels (A) and (B) use the same color code. For (A) and (B), statistical analyses were performed separately for each light intensity using one-way ANOVA followed by Tukey’s post-hoc test; different letters indicate significant differences (*p* < 0.05). **C)** Immunoblot of total protein extracts from plants grown under SD at LL, harvested 1 h after lights-on (+1 h), probed with a PEPC1-specific antibody. 50 µg of total protein extract was loaded in each well and 7 µl of molecular weight marker (Thermo ScientificTM PageRuler prestainedTM). MWM: Molecular Weight Marker; kDa: kiloDaltons.

Overall, proteome abundance, enzyme activity, and immunoblot validation support PEPC1 as a plausible contributor to the stronger phenotype observed in the *mdh1.1xme2* lines carrying the duplicated region.

### Low light promotes ammonium accumulation and shifts carbon/nitrogen balance in mdh1.1xme2 mutants

Because PEPC1 is a key integrator of carbon (C) and nitrogen (N) metabolism in leaves (Shi et al., 2015), we assessed whether the enhanced *mdh1.1xme2* phenotype is accompanied by altered elemental composition. We quantified total C and N in SD grown plants under LL and NL across the dial cycle. Under LL, the C/N ratio was consistently reduced in the double mutants, with the most pronounced differences detected at the end of the day and the end of the night (Suppl. Fig. 3A). This shift reflected both a decrease in total C throughout the diel cycle and an increase in total N at the end of the day (Suppl. Fig. 3A). In contrast, under NL we observed no significant genotype-dependent changes in C/N ratio, total C, or total N (Suppl. Fig. 3B).

To further examine N status, we quantified leaf ammonium (NH_4_^+^). Consistent with the bulk elemental measurements, NH_4_^+^ levels were indistinguishable among genotypes under NL, whereas under LL the *mdh1.1xme2* double mutants accumulated significantly higher NH_4_^+^ (Suppl. Fig. 3). Elevated NH_4_^+^ can restrict growth, impair chlorophyll accumulation, and induce photoprotective responses (Britto and Kronzucker, 2002; Podgórska et al., 2020), which is consistent with the pale-leaf and high-NPQ phenotypes observed in the double mutants.

### Light-regime-dependent accumulation of OAA-derived amino acids in the double mutants

To extend our analysis of the altered C/N balance and NH_4_^+^ accumulation observed in the double mutants with elevated PEPC1 activity, we next examined broader effects on central metabolism by untargeted GC-MS profiling of leaf extracts from plants grown under SD-LL, SD-NL, or LD-NL conditions. Samples were collected at three diel time points: 1 h before dawn (-1), 1 h after dawn (+1), and 1 h before the end of the photoperiod (+7/+15).

Principal component analysis (PCA) showed that samples were separated primarily by growth regime and sampling time, while also revealing a clear genotype-dependent shift, particularly under low light (Fig. 9). Under SD-LL, both mutant lines, *mdh1.1xme2.1* and *mdh1.1xme2.2*, clustered distinctly from Col-0, indicating a pronounced alteration of the metabolic profile under low irradiance. Separation between the -1 h and +1 h time points was additionally evident along PC2, consistent with a strong diel component. Under SD-NL, PC1 explained a larger fraction of the variance, and samples still showed genotype-associated clustering; however, the distance between *mdh1.1xme2* and Col-0 was reduced compared with SD-LL, suggesting partial normalization of the mutant metabolic profile under standard light. In LD-NL, sample separation was driven mainly by time after dawn, with the later time point (+15 h) displaced along PC1, whereas Col-0 and *mdh1.1xme2* overlapped more strongly at -1 h and +1 h.

**Figure 9.**
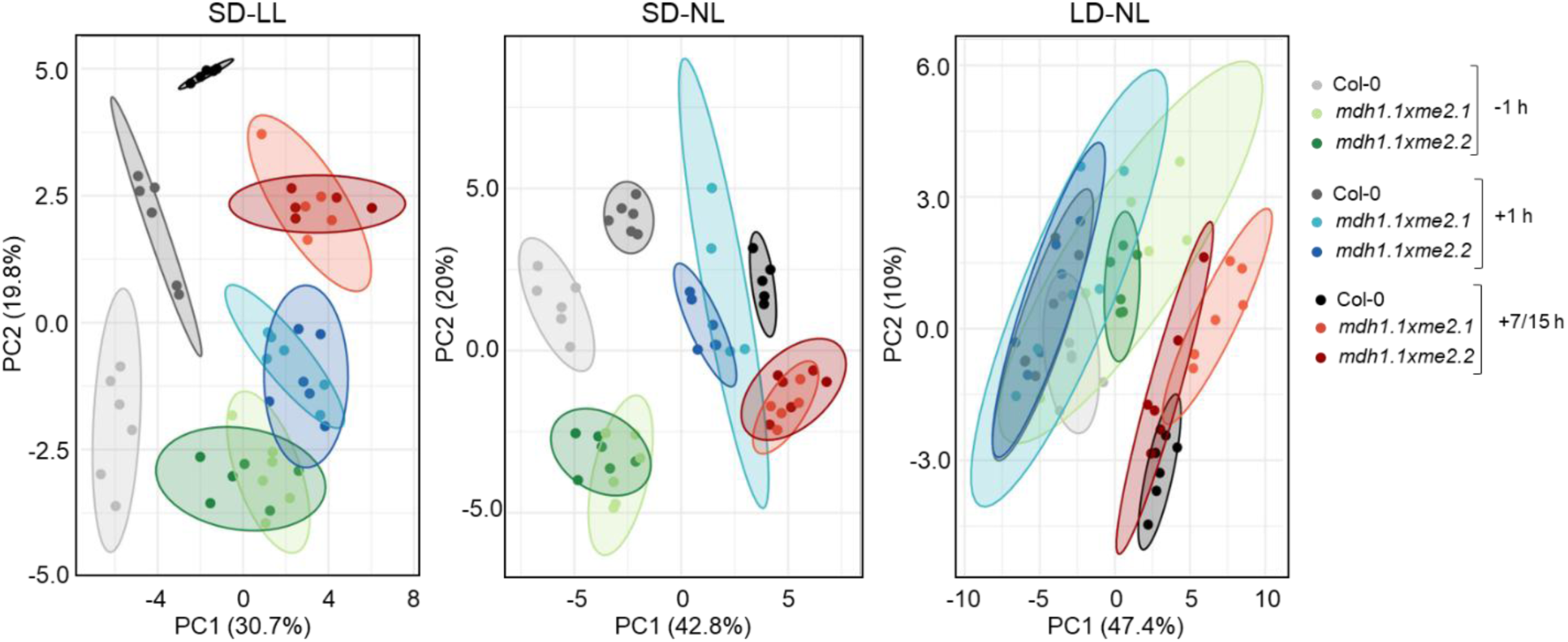
Principal component analysis (PCA) of metabolic profile from *mdh1.1xme2*. Metabolites measured by GC-MS from whole rosette grown under SD-LL, SD-NL and LD-NL harvested in three time points: one hour before illumination (-1 h), one hour after illumination (+1 h) and one hour before the end of the photoperiod (+7/15 h). Confidence ellipse of 80 %. Each dot represents one biological replicate.

Taken together, the PCA indicates that SD-LL enhances the *mdh1.1xme2*-specific shift in metabolic composition, whereas under NL conditions the genotype effect is weaker and more strongly shaped by diel progression.

In plants grown under SD-LL, intermediates of the tricarboxylic acid (TCA) cycle and related metabolites were substantially altered (Suppl. Fig. 4 and 5). Malate accumulated in the double mutant at +1 h and +7 h, whereas pyruvate was consistently reduced across all time points. The pyruvate-derived amino acids: alanine, valine, and leucine increased at the end of the night (-1 h). During the light period, valine remained elevated, while alanine declined significantly by +7 h, indicating differential regulation within this branch despite the overall reduction in pyruvate.

GC-MS profiling further revealed a pronounced and coordinated accumulation of oxaloacetate (OAA)-derived amino acids in the double mutant under SD-LL. Aspartate, lysine, threonine, isoleucine, and most prominently asparagine were elevated at all sampled time points (Suppl. Fig. 4 and 5; Fig. 10). While lysine, threonine, and isoleucine increased by approximately ∼4-times relative to wild type, asparagine exhibited an exceptional 15 to 20-fold rise. This dominant increase in asparagine is consistent with enhanced N assimilation and storage/transport capacity, aligning with the NH_4_^+^ accumulation and reduced C/N ratio observed under LL. Notably, this coordinated build-up of OAA-derived amino acids was not detected in *mdh1.1xme1xme2* grown under the same conditions (Suppl. Fig. 4 and 5; Fig. 10), indicating that the metabolic signature is specific to the duplication-associated double mutant. The concerted elevation of the aspartate family strongly suggests increased OAA availability and/or increased flux into OAA-dependent amino acid biosynthesis under LL. Given the increased PEPC abundance and activity in the double mutant, a parsimonious interpretation is that PEPC-dependent anaplerosis enhances OAA supply, which under low-energy conditions is preferentially diverted toward N-rich end products. In this framework, asparagine acts as a major nitrogen sink, potentially buffering excess reduced N when C availability is limited.

**Figure 10.**
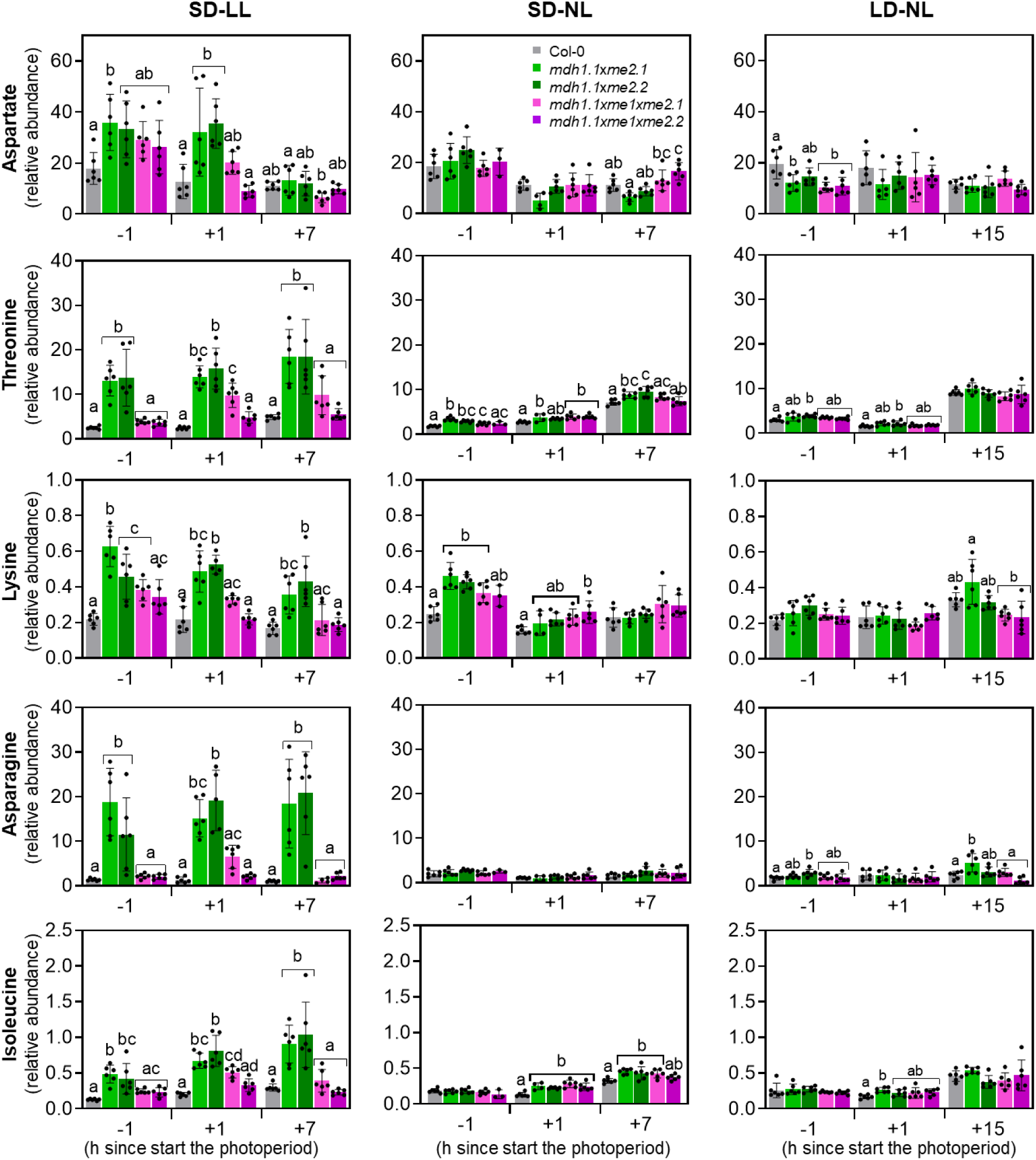
OAA-derived amino acids accumulate in *mdh1.1xme2* and *mdh1.1xme1xme2*. Relative metabolite abundances were determined by GC-MS in rosette leaves of the indicated genotypes grown under low light (LL) and normal light (NL) in short-day (SD) and long-day (LD) conditions. Samples were harvested 1 h before lights-on (-1), 1 h after lights-on (+1), and 1 h before lights-off (+7 /+15). Bars represent means ± SD (n = 6 independent biological replicates). For each time point, statistical significance was assessed separately using one-way ANOVA followed by Tukey’s post-hoc test; significant differences relative to Col-0 (*p* < 0.05) are indicated by asterisks. Data for the triple mutants were taken from Martinez et al. (2026).

Finally, the double mutants also showed increased accumulation of raffinose and proline (Suppl. Fig. 4 and 5), two metabolites commonly associated with low-light and carbon-limitation acclimation, as well as with stress-protective functions (Nishizawa-Yokoi et al., 2008; Szabados and Savouré, 2010; Verslues and Sharma, 2010; Jänkänpää et al., 2012). Their increase is consistent with a broader metabolic shift toward protective/maintenance states under LL, accompanying the diversion of C skeletons into N storage compounds.

## Conclusions and implications

### The *mdh1.1xme2* phenotype cannot be explained by NAD-ME deficiency alone

A central finding of this study is that the unexpectedly severe and light-regime-dependent phenotype of the *mdh1.1xme2* double mutants cannot be explained by the loss of MDH1 and ME2 alone. Based on genetic logic, *mdh1.1xme2* would be expected to resemble *mdh1.1xme1xme2* triple mutant if impaired heterodimeric NAD-ME capacity were the primary determinant. However, the double mutants were consistently smaller and exhibited stronger photosynthetic impairment, particularly under SD-LL. Importantly, multiple independent readouts demonstrated that ME1 protein and NAD-ME activity are undetectable in both double and triple mutant backgrounds, excluding residual NAD-ME1 function as an explanation for the phenotypic discrepancy.

### Genomic structural variation explains the unexpected phenotype

Whole-genome long-read sequencing resolved this apparent paradox. Both independent *mdh1.1xme2* lines carry a large ∼137-kb inverted duplication downstream of the *MDH1* T-DNA insertion, encompassing 38 annotated genes. This structural variant is absent from the *mdh1.1xme1xme2* triple mutant. The phenotype of the double-mutant therefore reflects a composite genotype: disruption of mitochondrial malate metabolism combined with increased dosage of dozens of additional genes. Consequently, interpreting the phenotype solely in terms of the intended loss-of-function alleles would be misleading.

Because our selection strategy tracked only the presence of the *MDH1* T-DNA insertion, meaning that the triple mutants could have been assembled from an insertion allele lacking the duplication, whereas the double-mutant lines likely derive from a second insertion-positive allele that carries the rearrangement.

### Gene dosage effects reshape metabolism

The duplication is functionally active rather than a silent passenger. RNA-seq revealed elevated transcript abundance across the duplicated interval, with the strongest coordinated effects around the night-to-day transition and under low irradiance.

Proteomics detected only a subset of proteins encoded in the interval, but among these, PEPC1 showed the most consistent increase in abundance. This observation was supported by higher PEPC activity, particularly under SD-LL conditions, and by immunoblot validation. Together, these findings identify PEPC1 as a plausible mechanistic link between the structural variant and downstream metabolic changes. Consistent with elevated PEPC capacity, the double mutants under SD-LL displayed a marked shift in C/N balance, including ammonium accumulation and a coordinated rise in OAA-derived (aspartate-family) amino acids, dominated by a strong increase in asparagine. This pattern suggests enhanced nitrogen storage and partitioning under carbon- and energy-limited conditions. In parallel, metabolites associated with stress tolerance and maintenance metabolism (e.g., raffinose and proline) accumulated, supporting a model in which low light amplifies duplication-driven dosage effects, redirecting metabolism toward nitrogen-rich sinks and protective acclimation states.

### Multiple candidate drivers within the duplicated interval

Although PEPC1 emerges as a compelling candidate contributor, the duplication likely affects phenotype interpretation in multiple ways. The interval contains several additional genes involved in signaling, proteostasis, and stress responses, any of which could influence growth, hormone homeostasis, chloroplast remodeling, or photosynthetic performance in a condition-dependent manner. Moreover, the inverted configuration and breakpoint context may alter the surrounding regulatory architecture. As a result, the observed expression changes may not follow a simple “two copies → twofold expression” relationship. PEPC1 should therefore be considered a strong candidate driver, but not yet the sole causal factor.

### Future directions and implications for functional genomics

Looking forward, disentangling “loss-of-function” from “duplication-driven” effects will require targeted genetic approaches. Priorities include: (i) generating clean *mdh1.1* knockouts without genomic structural rearrangements; (ii) backcrossing and segregating the duplication away from the *mdh1.1xme2* background, and (iii) testing whether increased PEPC1 dosage or activity is sufficient to reproduce key metabolic signatures in otherwise clean genetic backgrounds.

More broadly, our findings highlight an important methodological consideration for plant functional genomics. When mutant phenotypes deviate from genetic expectations, particularly in environment-dependent contexts, structural validation of insertion alleles becomes essential to ensure correct attribution of causality. In such cases, long-read genotyping or sequencing should be considered a critical step for fully resolving the underlying genotype and avoiding misinterpretation of mutant phenotypes (Pucker et al., 2021).

## Material and Methods

### Plant material

The plant lines used in this study include *Arabidopsis thaliana* L. Heynh. ecotype Columbia (Col-0, wild type), along with homozygous T-DNA insertion lines Sail-291-C05 (*me2.1*), SALK_131720 (*me2.2*; Tronconi et al., 2008), and GK-097C10 (*mdh1.1*; Tomaz et al., 2010) (Suppl. Table 1). The T-DNA insertion sites were verified by sequencing using specific primers (Suppl. Table 2; Suppl. Fig 1A). Double homozygous mutants were generated by genetic crosses of *mdh1.1* and *me2.1 or me2.2*. Homozygosity was confirmed by PCR using gene-specific and T-DNA left border primers (Suppl. Table 2; Suppl. Fig 1A).

### Plant growth

Plants were grown in soil (Floradur ® Substrate Multiplication Seed B, Rüttger Abstohs Gärtner-Einkauf GmbH&Co.KG, Siegburg, Germany) at 22/19 °C (day/night) under either long-day (LD: 16 h light/8 h dark) or short-day (SD: 8 h light/16 h dark) photoperiods, with light intensities of 60 µmol photons m^-2^ s^-1^ (LL) or 120 µmol photons m^-2^ s^-1^ (NL) and relative humidity of 70 %. Illumination was provided by a combination of Osram HO 80W/840 and HO 80W/830 bulbs. Seeds were stratified at 4 °C for 3 days prior to transfer to growth conditions.

For all the experiments, whole rosettes were harvested in SD at LL light intensity at 40 days after stratification (DAS), at NL at 30 DAS, and in LD at 23 days after stratification. If this set up does not apply in a given method, it is specified in its own section.

### RNA extraction and cDNA synthesis

Total RNA was extracted from rosette leaves of plants grown in SD under LL and NL harvested 40 DAS and 30 DAS, respectively, using the Universal RNA Purification Kit (Roboklon GmbH, Germany) according to the manufacturer’s instructions, and eluted in 30 µl of RNase-free distilled water. The harvest points were one hour before the light was on (-1), one hour after the light was on (+1) and one hour before the light was off (+7). RNA concentration and purity were determined using a NanoDrop Nabi™ UV/Vis spectrophotometer (MicroDigital Co. Ltd, South Korea), and RNA integrity was verified by agarose gel electrophoresis. Samples were stored at -20 °C until further use. To remove potential genomic DNA contamination, RNA samples were treated with the Ambion DNA-free™ DNA Removal Kit (Thermo Fisher Scientific, USA). Subsequently, 2 µg of RNA was reverse-transcribed into complementary DNA (cDNA) using RevertAid Reverse Transcriptase (Thermo Scientific, USA) and Oligo(dT) primers, following the manufacturer’s protocol.

### Reverse-transcription quantitative PCR

Reverse transcription quantitative PCR (RT-qPCR) was conducted using the KAPA SYBR FAST qPCR Kit (KAPA Biosystems Inc., Switzerland) to quantify the expression levels of *MDH1* and *ME2* in cDNA samples derived from the different plant lines. Reactions were performed on a Mastercycler® ep realplex system (Eppendorf, Germany), and cycle threshold (Ct) values were obtained using the accompanying realplex software. Gene expression was normalized to the *ACTIN2* (*ACT2I,* At3g18780) gene, which served as an internal reference.

### Leaf surface area

For the calculation of the leaf surface area and maximum diameter a sample size of n = 10 was used from each genotype. The growth process was visually documented throughout the plant development and under NL and LL in SD conditions. Each plant picture was then analyzed using the software ImageJ build 1.50e. The leaf surface areas of all plant lines were determinate using Rosette Tracker image analysis (De Vylder et al., 2012) and were then compared with each other using a two-way ANOVA followed by Tukey’s post-hoc test. Statistical analyses were performed with the software RStudio using the package “multicomp” (version:1.4-29; Hothorn et al., 2008) which compared the leaf surface area of each plant line with each time point (DAS), and the same was applied for the maximum diameter.

### Pulse amplitude modulation fluorometry

PAM (Pulse Amplitude Modulation) measurements were performed using the IMAGING-PAM Maxi version (Walz, Germany) with a PAR (Photosynthetically Active Radiation) of 81 µmol m^-2^ s^-1^. Plants had been dark-adapted for 20 min prior to each measurement. For each plant line, four biological replicates were analyzed, and five measurement points were selected from each replicate. Average values of five points from each biological replicate were used to calculate each photosynthetic parameter. The maximum quantum yield efficiency of PSII (Fv/Fm = [Fm-F0]/Fm) was taken as a measure of PSII intactness, quantum Yield of PSII (ΦPSII) indicates the amount of solar energy converted to chemical energy to be used by the plant cell, the electron transport rate (ETR) shows the speed of the electrons in the electron transport chain and the non-photochemical quenching (NPQ).

### Transmission electron microscopy

The fourth and fifth leaves from plants grown under SD conditions at 60 µE were excised one hour before the onset of the light period and immediately fixed in a solution containing 75 mM cacodylic acid sodium buffer (Carl Roth GmbH, Germany) pH 7 using HCl, 2.5% glutaraldehyde, 0.1% sucrose, and 2 mM MgSO_4_ for 4-5 hours at 4 °C in a desiccator. Following fixation, leaves were washed three times for 15 minutes each in 100 mM cacodylate buffer. Post-fixation was performed in 2% osmium tetroxide (OsO_4_) for 2.5 hours, followed by three 10-minute washes in distilled water. Leaves were then dehydrated through a graded acetone series (15%-50%), with three 10-minute incubations at each concentration. Subsequently, samples were incubated for 1 hour in 75% acetone containing 1% phosphotungstic acid and 1% uranyl acetate, followed by three 15-minute incubations in 80%, 90%, and 100% acetone. After dehydration, samples were infiltrated overnight with a 1:3 mixture of acetone and ERL-4206 resin (10% vinyl cyclohexene dioxide). This was followed by 4.5 hours in a 1:1 acetone/ERL mixture, 4 hours in a 3:1 acetone/ERL mixture, and finally overnight in pure ERL resin at 70 °C.

Once embedded, leaf sections were cut using an ultramicrotome and mounted on copper mesh grids. For contrast enhancement, grids were stained for 5 minutes in 1% uranyl acetate, followed by 5 minutes in a solution of lead nitrate [Pb(NO_3_)_2_] and sodium citrate [Na_3_C_6_H_5_O_7_·2H_2_O], and finally treated with 1 M NaOH, with water washes between each step. Prepared grids were examined using a Zeiss EM 10 transmission electron microscope (1978 model). Ultrathin sections were stained with Azur II 1% (w/v), Methylene blue 1% (w/v) dissolved in sodium tetraborate 1% (w/v) and observed using fluorescence stereomicroscope with 40x magnification lens (Leica MZFLIII).

### Determination of the phytohormone levels

Phytohormones were extracted from the whole rosette using 2-phase-extraction with isopropanol/H_2_O/methylene chloride as described previously (Pan et al., 2008; Wewer et al., 2026). Samples were harvested one hour before the light was on and frozen immediately in liquid nitrogen from plants grown under SD in LL and NL conditions. Tissue was ground using mortar and pestle in liquid nitrogen and approximately 50 mg of fresh tissue was used for each extraction. Phytohormone extracts were then analyzed by LC-MS. Phytohormone separation of phytohormones measured in negative ionization mode (*cis*,*trans*-ABA, JA and SA) was done on a C18 XSelect™ HSST3 HPLC column (2.5 µm, 3.0 mm x 150 mm (Waters)) at a flow rate of 0.35 mL/min using a binary gradient, where solvent A was H_2_O + 0.1 % (v/v) formic acid, and solvent B was methanol + 0.1 % (v/v) formic acid. The gradient was performed as described in (Wewer et al., 2026). Phytohormone separation of phytohormones measured in positive ionization mode (IAA, tZ9G, cZ9G, tZOG and iPR) was done on a C18 Atlantis™ Premier BEH column (2.5 µm, 2.1 mm X 150 mm (Waters)), with the same gradient. Phytohormone analysis by LC-MS was performed on a maXis 4G Q-TOF instrument (Bruker Daltonics), and quantification was based on relation of peak areas to stable isotope-labelled internal standards (I.S.) as described (Wewer et al., 2026). For comparing the different genotypes, statistical analysis was done using using a two-way ANOVA followed by Tukey’s post-hoc test using multcomp package (version:1.4-29; Hothorn et al., 2008) in R studio. *p*-val < 0.05 between the genotypes are considered significant differences.

### Determination of C/N ratio, ammonium concentration and metabolite profiling analysis

For the determination of C/N ratio, whole rosettes were harvested one hour before the end of the night (-1), one hour after (+1) and seven hours after (+7) the light was switched on from plants growing under LL and under NL in SD. For ammonium quantification, whole rosettes were harvested from plants grown under low light (LL) and normal light (NL) in short-day (SD) conditions, either 1 h before the onset of light or 1 h after lights-on. For metabolic profile analysis, the whole rosette leaves were harvested at three time points, one hour before the light was on (-1), one hour after light was on (+1) and one hour before the light was off (+7/+15) from plants grown under SD-LL, SD-NL and LD-NL and immediately frozen. Total carbon and nitrogen, ammonium content and metabolic profiles measured using gas chromatography-mass spectrometry (GC-MS) were determined according to Martinez et al. (2026). Statistical analyses were performed in RStudio using two-way ANOVA followed by Tukey’s post hoc test implemented in the multcomp package (version:1.4-29; Hothorn et al., 2008). Differences between genotypes were considered statistically significant at *p* < 0.05 (Supplementary Data 3).

### Isolation of mitochondria

Rosette from 8-week-old plants grown in SD at NL were harvested one hour after the light was turn on (+1). The isolation was done following an established protocol (Liu et al., 2023). Protein concentration was determinate using Bradford reactive (Thermo Fischer, Germany).

### NAD-ME activity assay

NAD-ME activity in isolated mitochondrial fractions was measured following the protocol described by Tronconi et al. (2008). Activity was assessed by monitoring NADH production using a Spark® microplate reader (Tecan) in 96-well UV-Star Microplates (Greiner Bio-One). The standard reaction mixture contained 50 mM MES-NaOH (pH 6.5), 4 mM NAD⁺, 10 mM L-malate, 5 mM dithiothreitol, 10 mM MnCl_2_, and 10 units of MDH from pig heart (Roche, Germany). Enzymatic activity was normalized to total protein content measured by Bradford (Thermo Fischer, Germany) in each sample.

### PEPC activity assay

PEPC activity was measured in protein extracts from plants grown under low light (LL) in short-day (SD) conditions. Whole rosettes were harvested 1 h after the onset of light (+1 h). Frozen tissue was ground twice for 15 s in a TissueLyser II (QIAGEN, Germany) at a frequency of 30 Hz. Three biological replicates were analyzed per genotype. Protein extraction was performed according to Hüdig et al. (2022), with slight modifications. Shortly, proteins were extracted by using 2 µl of extraction buffer (0.1 M HEPES pH 7.8, 0.1 M NaCl, 10 mM MgCl_2_, 0.5 % (v/v) Triton X-100, 10 mM DTT, 2 mM PMSF, 0.1% (w/v)) per mg of tissue. After the addition of the buffer samples were vortexed and incubated 20 min on ice. Afterwards, they were centrifugated at 10,000 g 4 °C 10 minutes. Supernatant was transferred to a pre-chilled tube and used for the PEPC activity determination. Protein concentration was accessed using Bradford reactive. PEPC activity was measured spectrophotometrically at 340 nm by coupling the production of oxaloacetate to the oxidation of NADH from malate dehydrogenase (MDH) at 30 °C in Spark® microplate reader (Tecan). The reaction mixture contained 50 mM HEPES pH 7.8, 10 mM MgCl_2_, 1 mM NaHCO_3_, 6 mM PEP, 0.15 mM NADH, and 10 U/ml MDH from pig heart (Roche, Germany). The crude extract was incubated with the reaction mixture without PEP till the signal at 340 nm was stabilized, after that the substrate was added to start the reaction and the absorbance was monitored for 20 min. Enzymatic activity defined by µmol of NADH consuming per min was normalized by the protein content in each extract.

### Immunoblot analysis

Immunoblot detection of ME1 was performed using mitochondria isolated from *A. thaliana* and a specific antibody raised against *A. thaliana* ME1 (ImmunoGlob GmbH, Germany). Recombinant *A. thaliana* ME1 was used as a positive control (Tronconi et al., 2008). Immunoblot detection of PEPC1 was performed using total protein extracts from rosettes and an antibody against *A. thaliana* PEPC1 (AS16 4110, Agrisera).

Denaturing polyacrylamide gel electrophoresis (SDS-PAGE) was carried out on 14% (w/v) polyacrylamide gels according to Laemmli (1970). Proteins were then transferred onto a nitrocellulose membrane for 1.5 h at 133 mA for ME1 and 200 mA for PEPC1 using a semi dry from Biorad system. Membranes were blocked for 1 h at room temperature in TBS (20 mM Tris-HCl, pH 7.5, 150 mM NaCl) supplemented with 5 % (w/v) milk powder. After blocking, the membrane was washed three times for 10 min using TBS-T (0.1 % (w/v) Tween 20 in TBS). For ME1 detection, the membrane was incubated overnight at 37 °C with 0.5 µg/µl of the primary antibody diluted in TBS containing 5 % (w/v) milk powder. For PEPC1 detection, the membrane was incubated overnight at 4 °C with the primary antibody diluted 1:1000 in TBS with 2 % (w/v) milk powder. Following primary antibody incubation, membranes were washed three times with TBS-T, following incubation with goat anti-rabbit HRP-conjugated secondary antibody, diluted 1:5000 for ME1 detection (Merck, Germany) or 1:40000 for PEPC1 detection (AS09602, Agrisera). After three additional washes with TBS-T, signal detection was performed by incubating the membranes for 5 min in a 1:1 mixture of ECL (enhanced chemiluminescence) substrate solution (Western Blot kit, Bio-Rad). Chemiluminescence signals were recorded using an Azure 300 imaging system (Azure Biosystems, USA) for ME1 and a ChemiDoc MP Imaging System (Bio-Rad) for PEPC1.

### Oxford nanopore technology sequencing

High molecular weight DNA was extracted from young *A. thaliana* plants using a modified CTAB-based method (Siadjeu et al., 2020). The Short Read Eliminator kit (Pacific Biosciences) was used to deplete short DNA fragments from the samples. Samples were prepared for nanopore sequencing following the SQK-LSK114 protocol starting with 1 µg of DNA. Sequencing was performed on R10.4.1 flow cells on a MinION Mk1B. POD5 files were subjected to basecalling with dorado v1.0.2 (https://github.com/nanoporetech/dorado) using the high accuracy model dna_r10.4.1_e8.2_400bps_hac@v5.2.0. Reads were aligned to the TAIR10 reference genome sequence with minimap v2.27-r1193 (Li, 2018) using default parameters for nanopore reads. Samtools v1.19.2 (Li et al., 2009) was used to convert SAM into BAM files for visualization in Integrative Genomics Viewer v2.7.2 (Robinson et al., 2011). Bedtools (Quinlan and Hall, 2010) was utilized to infer coverage for each genomic position (Pucker and Brockington, 2018). Loreta v1 (Pucker et al., 2021) was deployed for the investigation of T-DNA insertions and structural rearrangements. The sequences of pAC161 (AJ537514) and pCSA110 (McElver et al., 2001) served as bait for the discovery of T-DNA sequences. A dedicated Python scripts was applied for the genome-wide coverage analysis (available viaGitHub: https://github.com/bpucker/GKseq2).

### RNA seq analysis

For RNA extraction, rosette leaves were harvested one hour before and after the light was switched on from plants growing in SD under LL and NL. Total RNA was extracted with the SV Total RNA Isolation System (Promega) and genomic DNA was eliminated by treating the samples with DNase I using DNA-free™ Kit (Invitrogen, Thermo Fischer), 1 µl of rDNase I was used per sample incubating at 37 °C for 30 min. Total RNA was quantified by the Qubit RNA HS Assay (Thermo Fisher Scientific, MA, USA). The quality was assessed by capillary electrophoresis using the Fragment Analyzer and the ‘Total RNA Standard Sensitivity Assay’ (Agilent Technologies, Inc. Santa Clara, CA, USA). Three biological replicates per genotype and condition with RNA Quality Numbers (RQN) mean > 8.2 were analyzed. The library preparation was performed according to the manufacturer’s protocol using the ‘VAHTS™ Stranded mRNA-Seq Library Prep Kit’ for Illumina®. Shortly, 400 ng total RNA were used as input for mRNA capturing, fragmentation, the synthesis of cDNA, adapter ligation and library amplification. Bead purified libraries were normalized and finally sequenced on the NextSeq2000 system (Illumina Inc. San Diego, CA, USA) with a read setup of 1x100 bp. The BCL Convert Tool (version 3.8.4) was used to convert the BCL files to FASTQ files as well for adapter trimming and demultiplexing.

Data analyses on FASTQ files were conducted with CLC Genomics Workbench (version 23.0.2, Qiagen, Venlo, Netherlands). The reads of all probes were adapter trimmed (Illumina TruSeq) and quality trimmed (using the default parameters: bases below Q13 were trimmed from the end of the reads, ambiguous nucleotides maximal 2). Mapping was done against the *A. thaliana* (TAIR10.54) (July 20, 2022) genome sequence. The differential expression analysis was performed using the DEseq2 package (Love et al., 2014) from BiocManager in the software RStudio, gene with less than five reads were depreciated. The Resulting *p* values calculated by Wald test were corrected for multiple testing using Benjamini and Hochberg method (BH-adjusted *p* values). An adjusted *p* value of ≤ 0.05 was considered significant.

### Shotgun proteomic LC-MS analysis

Proteomic analysis was performed on plants grown under short-day (SD) conditions at light intensities of LL and NL. Whole rosettes were harvested 1.5 hours after the onset of light. Five biological replicates per genotype were collected. Protein extraction and digestion was carried out using a single-pot, solid-phase-enhanced sample preparation (SP3) protocol (Hughes et al., 2019), which was adapted to meet challenges inherent to plant samples (Mikulášek et al., 2021). Each extract was treated with 2 µg of pre-activated, sequencing grade Trypsin (V5111, Promega). Concentrations of extracted tryptic peptides were determined in a plate reader (Multiskan Sky, Thermo Fischer Scientific) using the Pierce Quantitative Colorimetric Peptide Assay Kit (Thermo Fisher Scientific, Dreieich, Germany) following the manufacturer’s instructions. Peptides were dried using a vacuum concentrator and resuspended in 0.1% (v/v) formic acid (FA) to yield a peptide concentration of 10 ng µl^-1^. Twenty microliters of the suspension were loaded on Evotips (Evosep) according to the manufacturer’s instructions for desalting and subsequent separation using an EvoSep One HPLC (Evosep). Peptides bound to the Evotip resin were eluted using the Whisper Zoom 40 SPD method and separated in a 15 cm Aurora Elite CSI column (inner diameter 75 µm, particle size 1.7 µm, pore size 120 Å, IonOpticks, Fitzroy, Australia) which was kept at a temperature of 50 °C. Eluting peptides were transferred to a timsTOF Pro mass spectrometer (Bruker, Germany) and analyzed using the pre-installed ‘DDA PASEF-short_gradient_0.5sec_cycletime’ method.

Acquired spectra were queried against an in-house modified TAIR10 database using the MaxQuant software (Cox and Mann, 2008) version 2.6.3.0. Default parameters were used, and the ‘iBAQ’ (Schwanhäusser et al., 2011) and ‘LFQ’ functions were additionally enabled for quantification of identified protein groups.

The resulting ‘proteingroups.txt’ file was complemented with predicted subcellular protein localization using the SUBAcon algorithm (Hooper et al., 2014) and Mapman annotation (Thimm et al., 2004). Clasified table was separated in two tables: plastid-protein groups and non-plastid-protein groups filtered by SUBAcon. These tables were submitted to statistical analysis by means of the Perseus software (Tyanova et al., 2016) version 2.1.2.0 in the form of two-sample t tests (*p* < 0.05). For this, protein groups matching ‘only identified by site’, ‘potential contaminant’ and ‘reverse’ were removed before LFQ values were log_2_-transformed. Protein groups with less than three values in each sample group were removed and missing values were imputed from normal distribution using a width of 0.3 and a down-shift of 1.8. A width normalization was performed volcano plots were produced using default parameters (FDR, 0.05; S_0_, 0.1).

### Data availability

The sequencing data generated for this study have been deposited in the European Nucleotide Archive (ENA) under accession number PRJEB96949 (https://www.ebi.ac.uk/ena/browser/view/PRJEB96949). The raw RNA-seq data (FASTQ files) supporting the findings of this study are available in the NCBI SRA repository, under BioProject accession PRJNA1427118 (https://www.ncbi.nlm.nih.gov/sra/PRJNA1427118). All MS raw files acquired in this study, along with the corresponding MaxQuant search results can be downloaded from the FTP server ftp://MSV000101094@massive-ftp.ucsd.edu.

## Author contributions

V.M. and M.P.M conceived the study. M.P.M., I.N., N.D., N.D., J.A.V.S.O., P.W., V.W. and H.E. performed experiments and collected data. V.M., M.P.M., B.P., H.E., I.F. and M.S. analyzed the data. V.M., M.P.M., I.F., and M.S. interpreted results. V.G.M and M.P.M. wrote the manuscript with input from all authors. V.M. supervised the project. V.M., I.F., and M.S. acquired funding.

## Acknowledgements

This work was supported by the Deutsche Forschungsgemeinschaft (DFG) through the collaborative research grant PAK918 (Project ID 289357231: MA2379/14-3 to VGM, FI 1655/3-3 to IF, and SCHW 1719/5-3 to MS), as part of the ‘Plant Mitochondria in New Light’ initiative. The Metabolism & Metabolomics Laboratories at the University of Cologne and Heinrich Heine University Düsseldorf are funded by the DFG under Germanýs Excellence Strategy, CEPLAS - EXC-2048/1 (project ID 390686111). Computational infrastructure and support for RNA-seq analyses were provided by the Centre for Information and Media Technology at Heinrich Heine University Düsseldorf, and additional support from the de.NBI Cloud within the German Network for Bioinformatics Infrastructure (de.NBI) and ELIXIR-DE at Forschungszentrum Jülich (W-de.NBI-001, W-de.NBI-004, W-de.NBI-008, W-de.NBI-010, W-de.NBI-013, W-de.NBI-014, W-de.NBI-016, W-de.NBI-022).

The co-authors have no conflict of interest to declare.

## REFERENCES

1. Akiyoshi DE, Klee H, Amasino RM, Nester EW, Gordon MP (1984) T-DNA of Agrobacterium tumefaciens encodes an enzyme of cytokinin biosynthesis. Proceedings of the National Academy of Sciences 81: 5994–5998

2. Alonso JM, Ecker JR (2006) Moving forward in reverse: genetic technologies to enable genome-wide phenomic screens in Arabidopsis. Nat Rev Genet 7: 524–536

3. Alonso JM, Stepanova AN, Leisse TJ, Kim CJ, Chen H, Shinn P, Stevenson DK, Zimmerman J, Barajas P, Cheuk R, et al (2003) Genome-Wide Insertional Mutagenesis of Arabidopsis thaliana. Science 301: 653–657

4. Azpiroz-Leehan R, Feldmann KA (1997) T-DNA insertion mutagenesis in *Arabidopsis*: going back and forth. Trends in Genetics 13: 152–156

5. Bashore C, Prakash S, Johnson MC, Conrad RJ, Kekessie IA, Scales SJ, Ishisoko N, Kleinheinz T, Liu PS, Popovych N, et al (2023) Targeted degradation via direct 26S proteasome recruitment. Nat Chem Biol 19: 55–63

6. Britto DT, Kronzucker HJ (2002) NH4+ toxicity in higher plants: a critical review. Journal of Plant Physiology 159: 567–584

7. Castle LA, Errampalli D, Atherton TL, Franzmann LH, Yoon ES, Meinke DW (1993) Genetic and molecular characterization of embryonic mutants identified following seed transformation in Arabidopsis. Molec Gen Genet 241: 504–514

8. Cheng H-P, Yang X-H, Lan L, Xie L-J, Chen C, Liu C, Chu J, Li Z-Y, Liu L, Zhang T-Q, et al (2020) Chemical Deprenylation of N6-Isopentenyladenosine (i6A) RNA. Angewandte Chemie International Edition 59: 10645–10650

9. Chilton M-D, Drummond MH, Merlo DJ, Sciaky D, Montoya AL, Gordon MP, Nester EW (1977) Stable incorporation of plasmid DNA into higher plant cells: the molecular basis of crown gall tumorigenesis. Cell 11: 263–271

10. Clark KA, Krysan PJ (2010) Chromosomal translocations are a common phenomenon in Arabidopsis thaliana T-DNA insertion lines. The Plant Journal 64: 990–1001

11. Cortleven A, Schmülling T (2015) Regulation of chloroplast development and function by cytokinin. J Exp Bot 66: 4999–5013

12. Cox J, Mann M (2008) MaxQuant enables high peptide identification rates, individualized p.p.b.-range mass accuracies and proteome-wide protein quantification. Nat Biotechnol 26: 1367–1372

13. De Vylder J, Vandenbussche F, Hu Y, Philips W, Van Der Straeten D (2012) Rosette Tracker: An Open Source Image Analysis Tool for Automatic Quantification of Genotype Effects. Plant Physiol 160: 1149–1159

14. Dello Ioio R, Linhares FS, Scacchi E, Casamitjana-Martinez E, Heidstra R, Costantino P, Sabatini S (2007) Cytokinins Determine *Arabidopsis* Root-Meristem Size by Controlling Cell Differentiation. Current Biology 17: 678–682

15. Feldmann KA (1991) T-DNA insertion mutagenesis in Arabidopsis: mutational spectrum. The Plant Journal 1: 71–82

16. de Framond AJ, Barton KA, Chilton M-D (1983) Mini–Ti: A New Vector Strategy for Plant Genetic Engineering. Nat Biotechnol 1: 262–269

17. Gagne JM, Downes BP, Shiu S-H, Durski AM, Vierstra RD (2002) The F-box subunit of the SCF E3 complex is encoded by a diverse superfamily of genes in Arabidopsis. Proceedings of the National Academy of Sciences 99: 11519–11524

18. Gelvin SB (2017) Integration of Agrobacterium T-DNA into the Plant Genome. Annual Review of Genetics 51: 195–217

19. Gelvin SB (2003) Agrobacterium-Mediated Plant Transformation: the Biology behind the “Gene-Jockeying” Tool. Microbiology and Molecular Biology Reviews 67: 16–37

20. Głowacka K, Kromdijk J, Leonelli L, Niyogi KK, Clemente TE, Long SP (2016) An evaluation of new and established methods to determine T-DNA copy number and homozygosity in transgenic plants. Plant Cell Environ 39: 908–917

21. Häusler RE, Geimer S, Kunz HH, Schmitz J, Dörmann P, Bell K, Hetfeld S, Guballa A, Flügge U-I (2009) Chlororespiration and Grana Hyperstacking: How an Arabidopsis Double Mutant Can Survive Despite Defects in Starch Biosynthesis and Daily Carbon Export from Chloroplasts. Plant Physiol 149: 515–533

22. Hoekema A, Hirsch PR, Hooykaas PJJ, Schilperoort RA (1983) A binary plant vector strategy based on separation of vir- and T-region of the Agrobacterium tumefaciens Ti-plasmid. Nature 303: 179–180

23. Holdorf MM, Owen HA, Lieber SR, Yuan L, Adams N, Dabney-Smith C, Makaroff CA (2012) Arabidopsis ETHE1 Encodes a Sulfur Dioxygenase That Is Essential for Embryo and Endosperm Development. Plant Physiol 160: 226–236

24. Hooper CM, Tanz SK, Castleden IR, Vacher MA, Small ID, Millar AH (2014) SUBAcon: a consensus algorithm for unifying the subcellular localization data of the Arabidopsis proteome. Bioinformatics 30: 3356–3364

25. Hothorn T, Bretz F, Westfall P (2008) Simultaneous Inference in General Parametric Models. Biometrical Journal 50: 346–363

26. Hüdig M, Tronconi MA, Zubimendi JP, Sage TL, Poschmann G, Bickel D, Gohlke H, Maurino VG (2022) Respiratory and C4-photosynthetic NAD-malic enzyme coexist in bundle sheath cell mitochondria and evolved via association of differentially adapted subunits. Plant Cell 34: 597–615

27. Hughes RA, Heron J, Sterne JAC, Tilling K (2019) Accounting for missing data in statistical analyses: multiple imputation is not always the answer. Int J Epidemiol 48: 1294–1304

28. Jänkänpää HJ, Mishra Y, Schröder WP, Jansson S (2012) Metabolic profiling reveals metabolic shifts in Arabidopsis plants grown under different light conditions. Plant, Cell & Environment 35: 1824–1836

29. Kleinboelting N, Huep G, Kloetgen A, Viehoever P, Weisshaar B (2012) GABI-Kat SimpleSearch: new features of the Arabidopsis thaliana T-DNA mutant database. Nucleic Acids Res 40: D1211–D1215

30. Koncz C, Mayerhofer R, Koncz-Kalman Z, Nawrath C, Reiss B, Redei GP, Schell J (1990) Isolation of a gene encoding a novel chloroplast protein by T-DNA tagging in Arabidopsis thaliana. EMBO J 9: 1337–1346

31. Krispil R, Tannenbaum M, Sarusi-Portuguez A, Loza O, Raskina O, Hakim O (2020) The Position and Complex Genomic Architecture of Plant T-DNA Insertions Revealed by 4SEE. International Journal of Molecular Sciences 21: 2373

32. Laemmli UK (1970) Cleavage of Structural Proteins during the Assembly of the Head of Bacteriophage T4. Nature 227: 680–685

33. Lee L-Y, Gelvin SB (2008) T-DNA Binary Vectors and Systems. Plant Physiol 146: 325–332

34. Li H (2018) Minimap2: pairwise alignment for nucleotide sequences. Bioinformatics 34: 3094–3100

35. Li H, Handsaker B, Wysoker A, Fennell T, Ruan J, Homer N, Marth G, Abecasis G, Durbin R (2009) The Sequence Alignment/Map format and SAMtools. Bioinformatics 25: 2078–2079

36. Liu Y-G, Mitsukawa N, Oosumi T, Whittier RF (1995) Efficient isolation and mapping of Arabidopsis thaliana T-DNA insert junctions by thermal asymmetric interlaced PCR. The Plant Journal 8: 457–463

37. Liu Y-T, Senkler J, Herrfurth C, Braun H-P, Feussner I (2023) Defining the lipidome of Arabidopsis leaf mitochondria: Specific lipid complement and biosynthesis capacity. Plant Physiol 191: 2185–2203

38. Love MI, Huber W, Anders S (2014) Moderated estimation of fold change and dispersion for RNA-seq data with DESeq2. Genome Biol 15: 550

39. Marks MD, Feldmann KA (1989) Trichome Development in Arabidopsis thaliana. I. T-DNA Tagging of the GLABROUS1 Gene. Plant Cell 1: 1043–1050

40. Martinez M del P, Nica I, Zheng K, Ditz N, Oliveira JAVS de, Barreto P, Blum N, Westhoff P, Pucker B, Eubel H, et al (2026) Mitochondrial malate metabolism acts as a control hub for photosynthesis and carbon-nitrogen balance in Arabidopsis. Biorxiv.

41. McElver J, Tzafrir I, Aux G, Rogers R, Ashby C, Smith K, Thomas C, Schetter A, Zhou Q, Cushman MA, et al (2001) Insertional Mutagenesis of Genes Required for Seed Development in Arabidopsis thaliana. Genetics 159: 1751–1763

42. Meinke DW, Meinke LK, Showalter TC, Schissel AM, Mueller LA, Tzafrir I (2003) A Sequence-Based Map of Arabidopsis Genes with Mutant Phenotypes,. Plant Physiol 131: 409–418

43. Mikulášek K, Konečná H, Potěšil D, Holánková R, Havliš J, Zdráhal Z (2021) SP3 Protocol for Proteomic Plant Sample Preparation Prior LC-MS/MS. Front Plant Sci 12: 635550

44. Nacry P, Camilleri C, Courtial B, Caboche M, Bouchez D (1998) Major Chromosomal Rearrangements Induced by T-DNA Transformation in Arabidopsis. Genetics 149: 641–650

45. Nicolia A, Ferradini N, Veronesi F, Rosellini D (2017) An Insight into T-DNA Integration Events in Medicago sativa. International Journal of Molecular Sciences 18: 1951

46. Nishizawa-Yokoi A, Yabuta Y, Shigeoka S (2008) The contribution of carbohydrates including raffinose family oligosaccharides and sugar alcohols to protection of plant cells from oxidative damage. Plant Signaling & Behavior 3: 1016–1018

47. Olive J, Vallon O (1991) Structural organization of the thylakoid membrane: freeze-fracture and immunocytochemical analysis. J Electron Microsc Tech 18: 360–374

48. O’Malley RC, Alonso JM, Kim CJ, Leisse TJ, Ecker JR (2007) An adapter ligation-mediated PCR method for high-throughput mapping of T-DNA inserts in the Arabidopsis genome. Nat Protoc 2: 2910–2917

49. O’Malley RC, Barragan CC, Ecker JR (2015) A User’s Guide to the Arabidopsis T-DNA Insertion Mutant Collections. *In* JM Alonso, AN Stepanova, eds, Plant Functional Genomics: Methods and Protocols. Springer, New York, NY, pp 323–342

50. Pan X, Welti R, Wang X (2008) Simultaneous quantification of major phytohormones and related compounds in crude plant extracts by liquid chromatography–electrospray tandem mass spectrometry. Phytochemistry 69: 1773–1781

51. Podgórska A, Mazur R, Ostaszewska-Bugajska M, Kryzheuskaya K, Dziewit K, Borysiuk K, Wdowiak A, Burian M, Rasmusson AG, Szal B (2020) Efficient Photosynthetic Functioning of Arabidopsis thaliana Through Electron Dissipation in Chloroplasts and Electron Export to Mitochondria Under Ammonium Nutrition. Front Plant Sci 11: 103

52. Pucker B, Brockington SF (2018) Genome-wide analyses supported by RNA-Seq reveal non-canonical splice sites in plant genomes. BMC Genomics 19: 980

53. Pucker B, Kleinbölting N, Weisshaar B (2021) Large scale genomic rearrangements in selected Arabidopsis thaliana T-DNA lines are caused by T-DNA insertion mutagenesis. BMC Genomics 22: 599

54. Quinlan AR, Hall IM (2010) BEDTools: a flexible suite of utilities for comparing genomic features. Bioinformatics 26: 841–842

55. Robinson JT, Thorvaldsdóttir H, Winckler W, Guttman M, Lander ES, Getz G, Mesirov JP (2011) Integrative genomics viewer. Nat Biotechnol 29: 24–26

56. Rosso MG, Li Y, Strizhov N, Reiss B, Dekker K, Weisshaar B (2003) An Arabidopsis thaliana T-DNA mutagenized population (GABI-Kat) for flanking sequence tag-based reverse genetics. Plant Mol Biol 53: 247–259

57. Roussin-Léveillée C, St-Amand M, Desbiens-Fortin P, Perreault R, Pelletier A, Gauthier S, Gaudreault-Lafleur F, Laforest-Lapointe I, Moffett P (2025) Co-occurrence of chloroplastic ROS production and salicylic acid induction in plant immunity. New Phytologist 248: 1989–2004

58. Schröder G, Waffenschmidt S, Weiler EW, Schröder J (1984) The T-region of Ti plasmids codes for an enzyme synthesizing indole-3-acetic acid. European Journal of Biochemistry 138: 387–391

59. Schwanhäusser B, Busse D, Li N, Dittmar G, Schuchhardt J, Wolf J, Chen W, Selbach M (2011) Global quantification of mammalian gene expression control. Nature 473: 337–342

60. Sessions A, Burke E, Presting G, Aux G, McElver J, Patton D, Dietrich B, Ho P, Bacwaden J, Ko C, et al (2002) A High-Throughput Arabidopsis Reverse Genetics System. Plant Cell 14: 2985–2994

61. Shi J, Yi K, Liu Y, Xie L, Zhou Z, Chen Y, Hu Z, Zheng T, Liu R, Chen Y, et al (2015) Phosphoenolpyruvate Carboxylase in Arabidopsis Leaves Plays a Crucial Role in Carbon and Nitrogen Metabolism. Plant Physiol 167: 671–681

62. Siadjeu C, Pucker B, Viehöver P, Albach DC, Weisshaar B (2020) High Contiguity de novo Genome Sequence Assembly of Trifoliate Yam (Dioscorea dumetorum) Using Long Read Sequencing. Genes (Basel) 11: 274

63. Sussman MR, Amasino RM, Young JC, Krysan PJ, Austin-Phillips S (2000) The Arabidopsis Knockout Facility at the University of Wisconsin–Madison1. Plant Physiol 124: 1465–1467

64. Szabados L, Savouré A (2010) Proline: a multifunctional amino acid. Trends in Plant Science 15: 89–97

65. Szarzanowicz MJ, Waldburger LM, Busche M, Geiselman GM, Kirkpatrick LD, Kehl AJ, Tahmin C, Kuo RC, McCauley J, Pannu H, et al (2025) Binary vector copy number engineering improves Agrobacterium-mediated transformation. Nat Biotechnol 43: 1708–1716

66. Tamura K, Kawabayashi T, Shikanai T, Hara-Nishimura I (2016) Decreased Expression of a Gene Caused by a T-DNA Insertion in an Adjacent Gene in Arabidopsis. PLOS ONE 11: e0147911

67. Tax FE, Vernon DM (2001) T-DNA-Associated Duplication/Translocations in Arabidopsis. Implications for Mutant Analysis and Functional Genomics. Plant Physiol 126: 1527–1538

68. Thimm O, Bläsing O, Gibon Y, Nagel A, Meyer S, Krüger P, Selbig J, Müller LA, Rhee SY, Stitt M (2004) mapman: a user-driven tool to display genomics data sets onto diagrams of metabolic pathways and other biological processes. The Plant Journal 37: 914–939

69. Thomas CM, Jones DA, English JJ, Carroll BJ, Bennetzen JL, Harrison K, Burbidge A, Bishop GJ, Jones JDG (1994) Analysis of the chromosomal distribution of transposon-carrying T-DNAs in tomato using the inverse polymerase chain reaction. Molec Gen Genet 242: 573–585

70. Tomaz T, Bagard M, Pracharoenwattana I, Lindén P, Lee CP, Carroll AJ, Ströher E, Smith SM, Gardeström P, Millar AH (2010) Mitochondrial malate dehydrogenase lowers leaf respiration and alters photorespiration and plant growth in Arabidopsis. Plant Physiol 154: 1143–1157

71. Tronconi MA, Fahnenstich H, Gerrard Weehler MC, Andreo CS, Flügge U-I, Drincovich MF, Maurino VG (2008) Arabidopsis NAD-Malic Enzyme Functions As a Homodimer and Heterodimer and Has a Major Impact on Nocturnal Metabolism. Plant Physiol 146: 1540–1552

72. Tyanova S, Temu T, Cox J (2016) The MaxQuant computational platform for mass spectrometry-based shotgun proteomics. Nat Protoc 11: 2301–2319

73. Valentine ME, Wolyniak MJ, Rutter MT (2012) Extensive Phenotypic Variation among Allelic T-DNA Inserts in Arabidopsis thaliana. PLOS ONE 7: e44981

74. Van Larebeke N, Genetello C, Schell J, Schilperoort RA, Hermans AK, Hernalsteens JP, Van Montagu M (1975) Acquisition of tumour-inducing ability by non-oncogenic agrobacteria as a result of plasmid transfer. Nature 255: 742–743

75. Verslues PE, Sharma S (2010) Proline metabolism and its implications for plant-environment interaction. Arabidopsis Book 8: e0140

76. Wang K, Herrera-Estrella L, Montagu MV, Zambryski P (1984) Right 25 by terminus sequence of the nopaline t-DNA is essential for and determines direction of DNA transfer from Agrobacterium to the plant genome. Cell 38: 455–462

77. Wewer V, Dyballa-Rukes N, Metzger S (2026) Quantification of Phytohormones in Plants - Optimized Extraction, Separation and Detection. Biorxiv.

78. van Wijk KJ, Kessler F (2017) Plastoglobuli: Plastid Microcompartments with Integrated Functions in Metabolism, Plastid Developmental Transitions, and Environmental Adaptation. Annu Rev Plant Biol 68: 253–289

79. Zambryski P, Tempe J, Schell J (1989) Transfer and function of T-DNA genes from Agrobacterium Ti and Ri plasmids in plants. Cell 56: 193–201

80. Zechmann B (2019) Ultrastructure of plastids serves as reliable abiotic and biotic stress marker. PLOS ONE 14: e0214811

