## Supplemental Figures for "A hidden T-DNA-linked inversion-duplication causes a pronounced light-dependent phenotype in Arabidopsis"

### Supplementary Figures

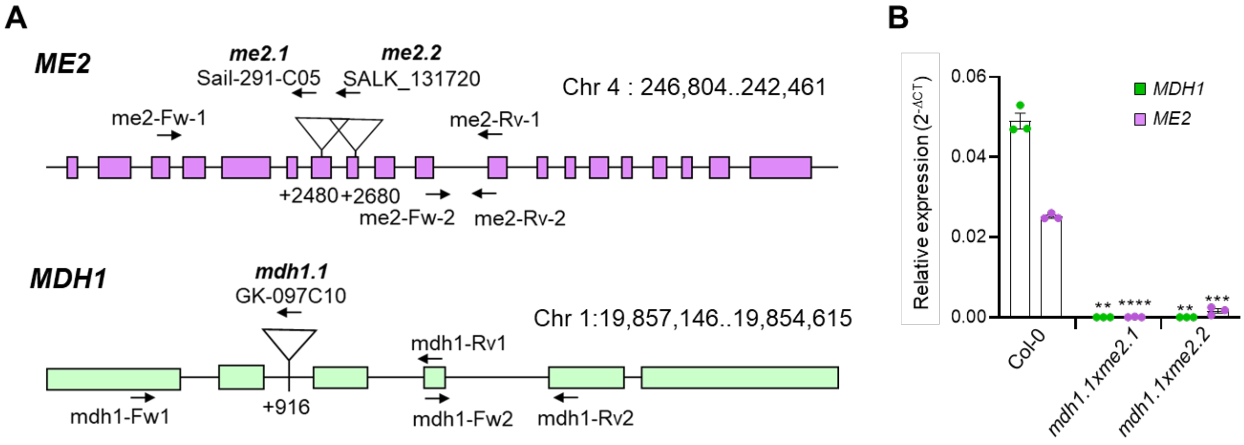


**Supplementary Figure 1. T-DNA insertion lines of *A. thaliana* mitochondrial *ME2* and *MDH1*. A)** Gene models of *ME2*and *MDH1* indicating the confirmed T-DNA insertion sites. Black arrows indicate locations of primer binding sites used for zygosity analysis and RT-qPCR (Suppl. Table 2). Rectangles represent exons and connecting lines represent introns. **B)** Relative expression levels of *ME2* and *MDH1* in rosette leaves of 40-day-old Col-0 and double insertional mutants grown in SD and LL and harvested 1 h before onset of the light. Data represent means ± SD (n = 3). Si Significance relative to Col-0 was assessed by Welch's t-test ($p < 0.05$, *; $p < 0.01$, **; $p < 0.001$, ***; $p < 0.0001$, ****).

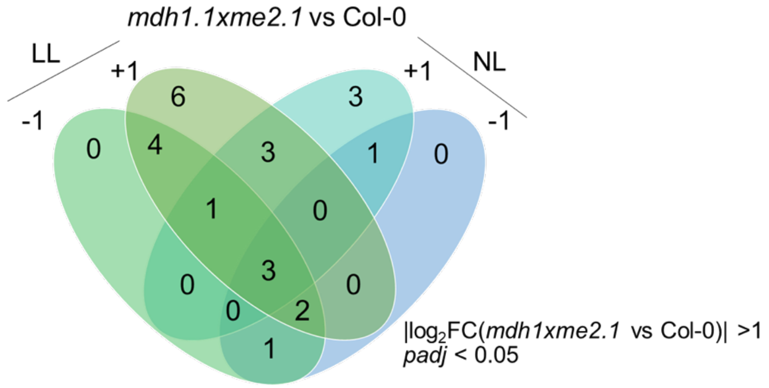


**Supplementary Figure 2. Analysis of the differential expressed genes from the duplicated region under the light-dark transition in SD.** Venn diagram (created using Venny 2.1) showing common and condition-unique high abundance transcripts from the duplicated region in *mdh1xme2.1* at LL and NL and -1 and +1 (high abundance transcripts: $log_2FC > 1$; n = 3 per genotype and condition; Supplementary Data 1). LL: low light; NL: normal light.

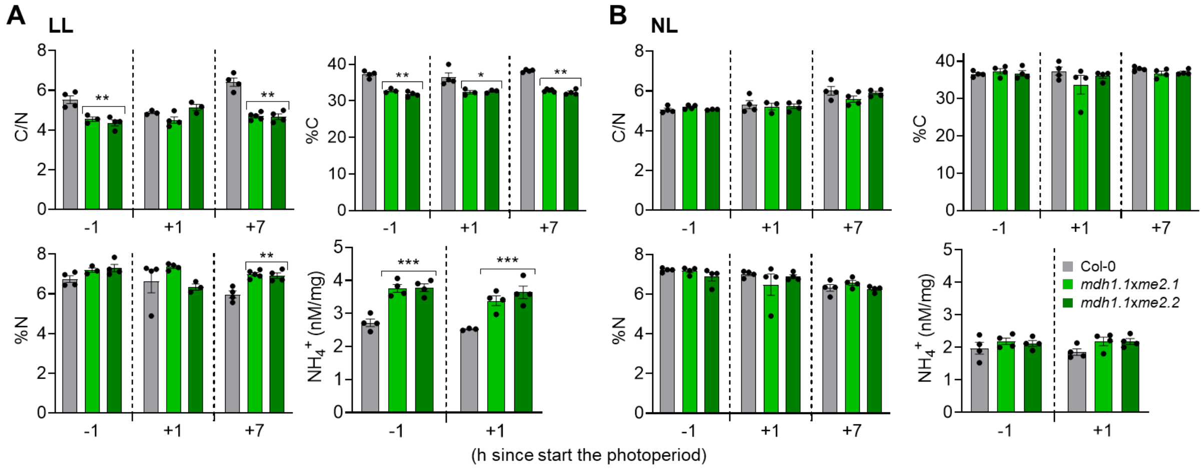


**Supplementary Figure 3. Carbon-to-nitrogen balance and ammonium accumulation.** Carbon-to-nitrogen ratio (C/N), calculated as total carbon divided by total nitrogen, total carbon (%C) and total nitrogen (%N) (percent of dry weight) were quantified in rosette leaf samples harvested 1 h before lights-on (-1 h), 1 h after lights-on (+1 h), and 1 h before lights-off (+7 h). Total leaf $NH_4^+$ concentrations were measured at -1 h and +1 h. Plants were grown under short-day (SD) conditions at low light (LL; **A**) or normal light (NL; **B**). Data represent means ± SD from 3-4 independent biological replicates per genotype. Significance relative to Col-0 was determined using Welch's t-test ($p < 0.05$, *; $p < 0.01$, **; $p < 0.001$, ***; $p < 0.0001$, ****).

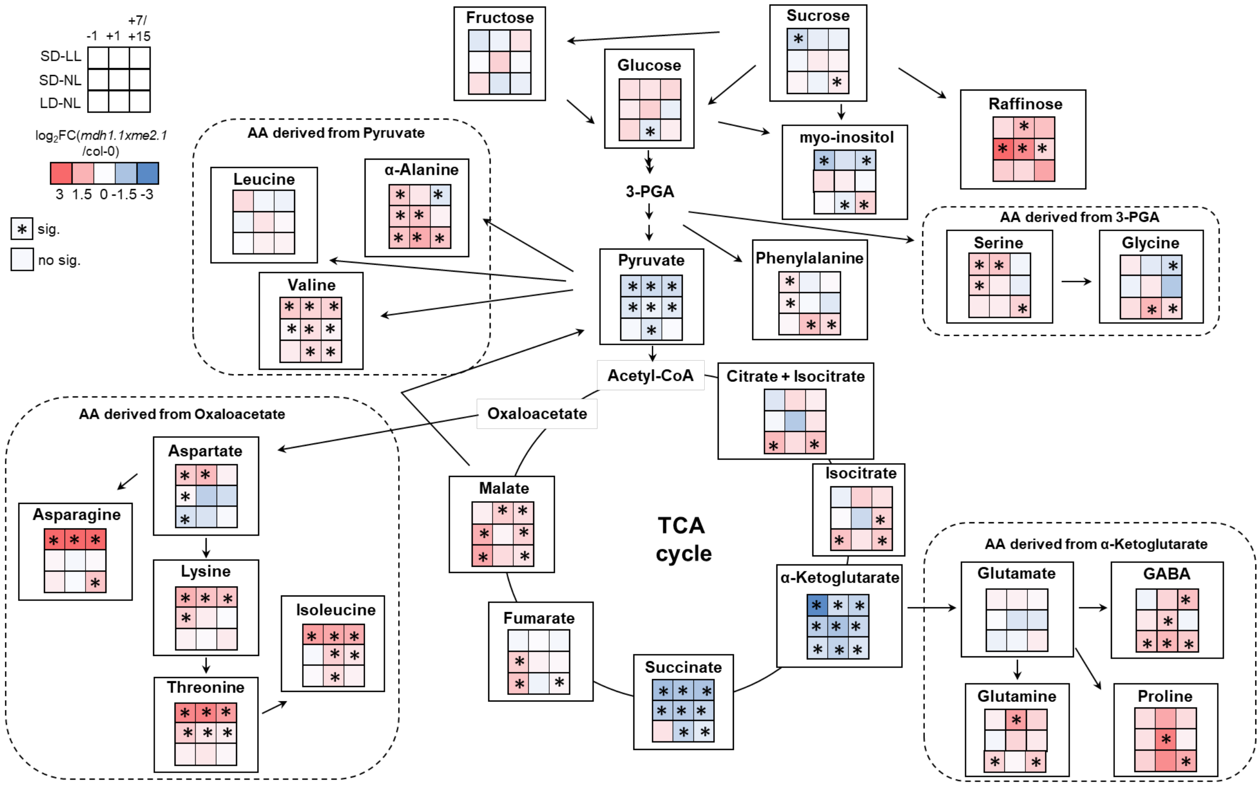


**Supplementary Figure 4. Relative metabolite abundance.** Rosette of *mdh1.1xme2.1* and Col-0 were harvested 1 h before lights-on (-1 h), 1 h after lights-on (+1 h) and 1 hour before lights-off (+7/15) grown in SD at LL and NL, and at NL in LD. Statistical analyses of n=6 independent biological replicates were performed for each time point using ANOVA followed by Tukey's HSD. Asterisks show metabolites that differ significantly at *p-val* < 0.05 from the wild type.

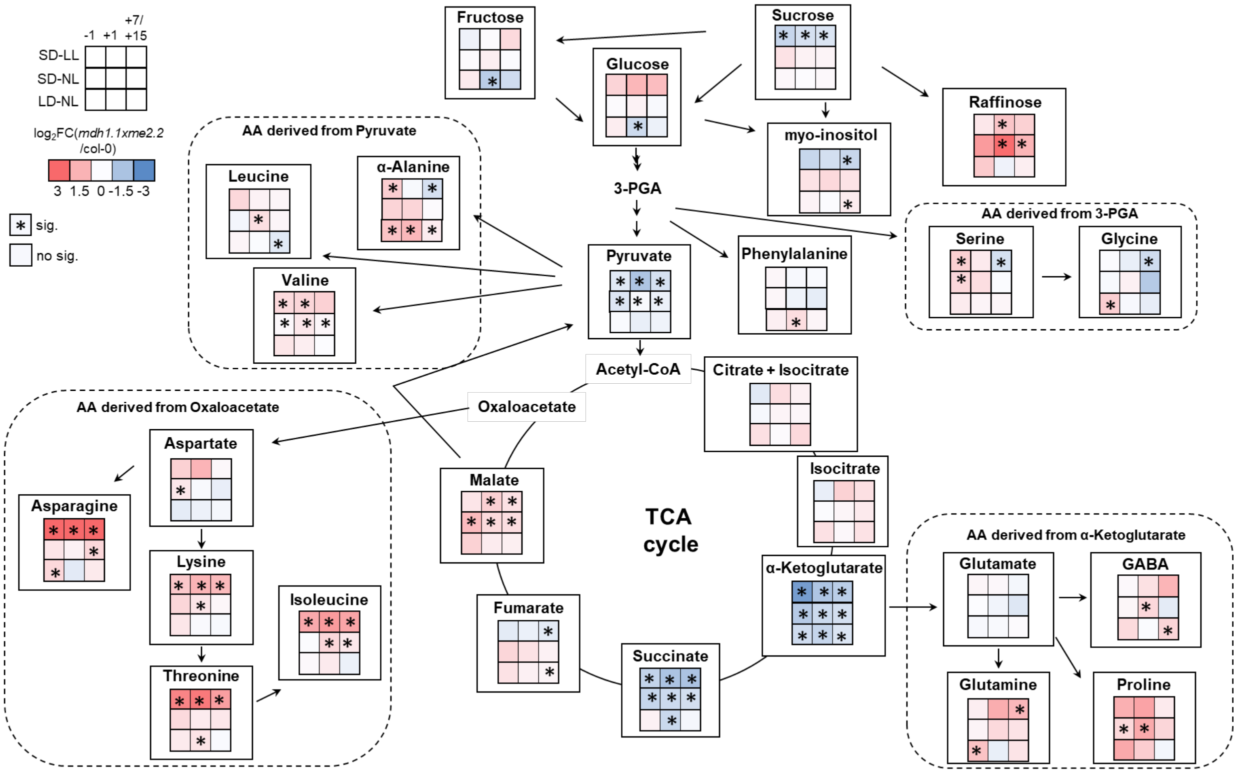


**Supplementary Figure 5. Relative metabolite abundance.** Rosette of *mdh1.1xme2.2* and Col-0 were harvested 1 h before lights-on (-1 h), 1 h after lights-on (+1 h) and 1 hour before lights-off (+7/15) grown in SD at LL and NL, and at NL in LD. Statistical analyses of n=6 independent biological replicates were performed for each time point using ANOVA followed by Tukey's HSD. Asterisks show metabolites that differ significantly at *pval* < 0.05 from the wild type.
