## Supplemental Tables for "A hidden T-DNA-linked inversion-duplication causes a pronounced light-dependent phenotype in Arabidopsis"

### Supplementary Tables

**Supplementary Table 1.** Information on the T-DNA insertion lines used in the work. s. a., see above; ko. knock out; kd, knock down.

| **Mutant (Type)** | **Gene** | **Gene locus** | **Insertion line** | **Reference** | **Phenotype SD-LL** |
|---|---|---|---|---|---|
| *mdh1.1* (ko) | *MDH1* | At1g53240 | GK-097C10 | (Tomaz et al., 2010) | no |
| *me1* (ko) | *ME1* | At2g13560 | Sail-374-A02 | (Tronconi et al., 2008) | no |
| *me2.1* (ko) | *ME2* | At4g00570 | Sail-291-C05 | | no |
| *me2.2* (kd) | *ME2* | At4g00570 | SALK_131720 | | no |
| *mdh1.1xme1xme2.1* | s.a. | s.a. | s.a. | (Martinez et al., 2026) | yes |
| *mdh1.1xme1xme2.2* | s.a. | s.a. | s.a. | | yes |
| *mdh1.1xme2.1* | s.a. | s.a. | s.a. | this work | yes |
| *mdh1.1xme2.2* | s.a. | s.a. | s.a. | | yes |

**Supplementary Table 2.** List of primers used in this study. s. a., see above.

| **Name** | **Sequence (5'- 3')** | **Gene/T-DNA Insertion** | **AGIs** |
|---|---|---|---|
| me2-Fw1 | GACCTGTGTACAGCAATGTGATCG | *ME2* | At4g00570 |
| me2-Rv1 | CCTGGACATCATCATTGAACATGC | | |
| mdh1-Fw1 | TCCGATCTTCTGCCTCC | *MDH1* | At1g53240 |
| mdh1-Rv1 | CAACTGGGACATTTGCCTT | | |
| act2-Fw1 | TAACTCTCCCGCTATGTATGTCGC | *ACT2* | At3g18780 |
| act2-Rv1 | GAAGCAAGAATGGAACCACCG | | |
| Sail | TAGCATCTGAATTTCATAACCAATCTCGAT ACAC | *ME2.1* | s. a. |
| SALK | TGGTTCACGTAGTGGGCCATCG | *ME2.2* | s. a. |
| GABI | ATATTGACCATCATACTCATTGC | *MDH1* | s. a. |
| me2-Fw2 | TTGGTGTAACTAAGATGGCAGT | *ME2* | s. a. |
| me2-Rv2 | CGTATCTCAGCTGGATTTTTGG | | |
| mdh1-Fw2 | TGTCCTCTCCGTGGTGACAAGACC | *MDH1* | s. a. |
| mdh1-Rv2 | TCCGATCTTCTGCCTCC | | |
| act2-Fw2 | CTTGCACCAAGCAGCATGAA | *ACT2* | s. a. |
| act2-Rv2 | CCGATCCAGACACTGTACTTCCTT | | |
